# Neuronal population dynamics during motor plan cancellation in non-human primates

**DOI:** 10.1101/774307

**Authors:** Pierpaolo Pani, Margherita Giamundo, Franco Giarrocco, Valentina Mione, Emiliano Brunamonti, Maurizio Mattia, Stefano Ferraina

**Author notes:** Corresponding author: Address correspondence to: Prof. Stefano Ferraina, MD PhD, Department of Physiology and Pharmacology P.le Aldo Moro 5 – CU027, 00185 Rome, Italy. These authors contributed equally (Maurizio Mattia, Stefano Ferraina).

## Abstract

To understand the cortical neuronal dynamics behind movement generation and control most studies focused on tasks where actions were planned and then executed, using different instances of visuomotor transformations. However, to fully understand the dynamics related to movement control one must also study how movements are actively inhibited. Inhibition, indeed, represents the first level of control both when different alternatives are available and only one solution could be adopted, and when it is necessary to maintain the current position.

We recorded neuronal activity from a multielectrode array in the dorsal premotor (PMd) cortex of monkeys performing a countermanding reaching task that requires, in a subset of trials, to cancel a planned movement before its onset. In the analysis of the neuronal state-space of PMd we found a subspace in which activities conveying temporal information were confined during active inhibition and position holding. Movement execution required activities to escape from this subspace toward an orthogonal subspace and, furthermore, surpass a threshold associated with the maturation of the motor plan. These results revealed further details in the neuronal dynamics underlying movement control, extending the hypothesis that neuronal computation confined in an ‘output-null’ subspace does not produce movements.

**Significance Statement:** A core question in neuroscience is how the brain generates arm movements. Most studies have approached this issue by investigating the neuronal dynamics that accompany movement production, leaving unanswered the question of which aspects of this dynamics are logically necessary to make the movement. Here we explored this topic by characterizing the neuronal correlates of movement decisions between active inhibition and release of movements. We found that active inhibition and stillness require neuronal signals to be confined in a functional sub-space while actions depend on the transit of activities in an orthogonal space. This dynamics is characterized by a threshold mechanism finally allowing the translation of the motor plan into overt action.

## Introduction

Our brain controls voluntary movements. However, the cortical computation behind this control capability is still under investigation.

To uncover how neurons in motor areas participate in arm movement control, most of the studies focused on different versions of the delayed reaching task. Here, a cue signal providing information about movement parameters is given ahead of the Go signal finally instructing to move. The delay epoch allows to easily separate signal-related activities from those more related to movement execution (1, 2). Using the delayed reaching task and different approaches, in the last 15 years the dorsal premotor cortex (PMd) has been suggested to be a key area in which, during motor preparation, a cascade of neuronal events bring the collective activity into a preferred and stereotyped (attractor-like) state (3, 4, 5). Such preparatory state (1) allows neuronal population activity to vary after sensory instructions without eliciting any movement onset, thus defining an ‘output-null’ state well separated by the inner representation of the successive movement execution (6, 7). Within this framework, movement generation is the consequence of a further change in the neuronal state (6, 7), corresponding to the transition from the ‘output-null’ towards the ‘output-potent’ space. However, all these studies exploited the delayed reaching task which always requires the generation of the planned movement. But what happens when the planned movement is cancelled? How is the neuronal ensemble dynamics of motor cortices reshaped in this case? Only comparing conditions in which a movement is performed *versus* those in which the same prepared movement is cancelled can reveal whether the unfolding of the specific neuronal dynamics is required for the movement to be made.

To tackle this issue, we recorded single neuronal activity from PMd of rhesus monkeys while performing the countermanding task (8–12). In this task, a Stop signal presented in some trials during the reaction time asks for the cancellation of a movement during the motor plan development. In the population of neurons simultaneously recorded, we applied a state-space approach to reveal whether a specific neuronal dynamics is necessary to generate a movement. Put it simply: if a neuronal dynamics underlies movement generation, it must occur when the movement is executed, while it must be absent when the movement is actively withheld. Here we propose and test a generalization of the hypothesis that motor-related neuronal activity must be confined within an ‘output-null’ state region, to successfully suppress the translation of motor plans into overt movements, and that only the pull-out from this region can lead to the movement. Such confinement is an active process determined by a possible ‘attractive’ capability of this state region, and the ‘escape’ from it in the network dynamics is found to be due to a specific contribution of a heterogeneous set of single units.

## Results

Monkeys (P and C) after completing their training period performed a countermanding reaching task with 66% of **no-stop trials** and 34% of **stop trials** (Fig. 1A), and targets located either to the right or to the left of the workspace. For the neuronal analysis we selected two (one for each animal) of those experimental sessions that had the highest number of trials and well isolated units, in which the animal behavior was compatible with the race model hypotheses (independence assumption; see Fig. S1A, B; Table S1, and Material and Methods) such that a reliable estimate of the **stop-signal reaction time (SSRT;** see Table S1) could be computed. This is because the SSRT is an estimate of the time necessary to suppress the movement in the task, a needed information to further identify the dynamical organization of PMd units participating in movement inhibition.

**Figure 1.**
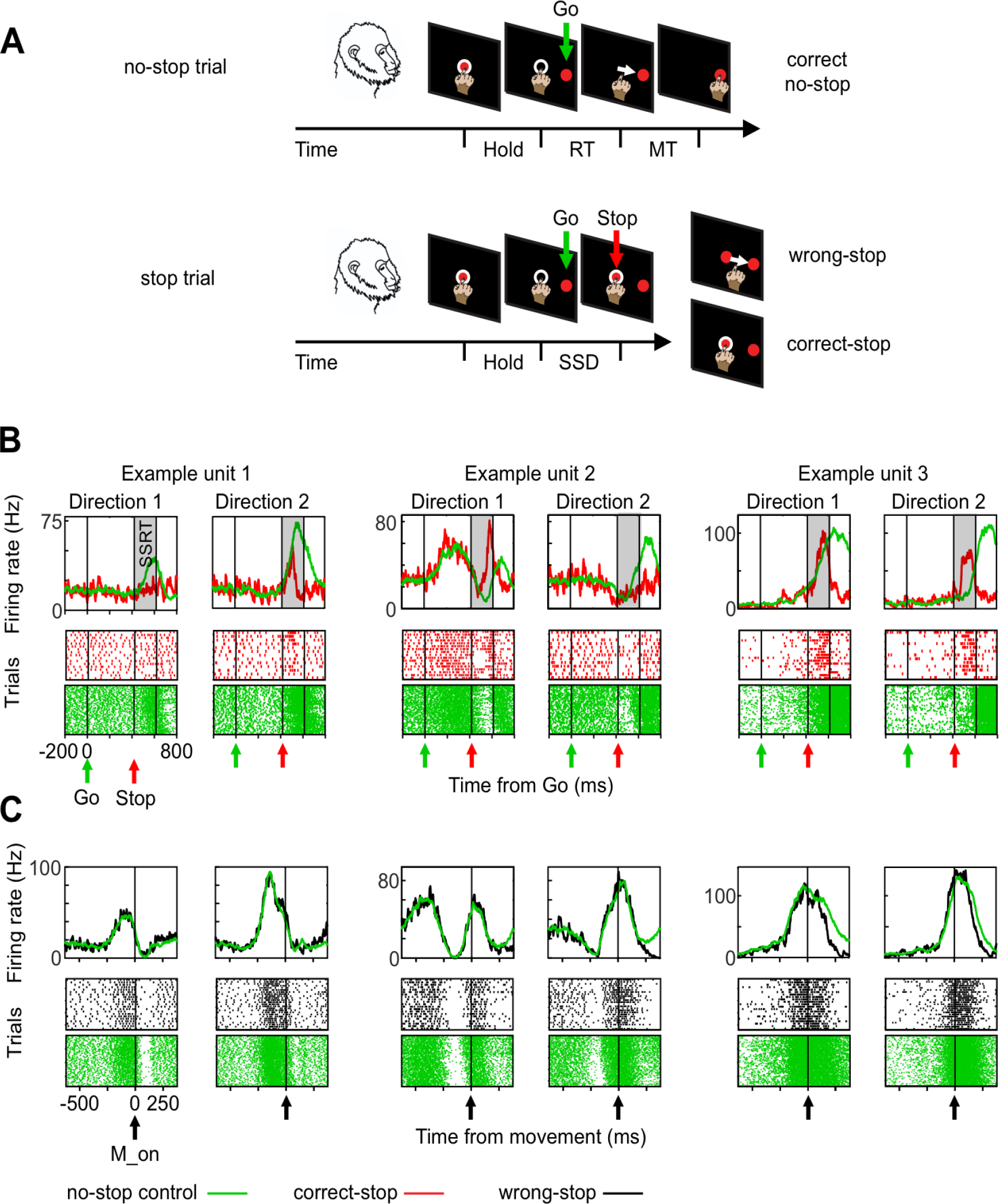
Behavioral task and heterogeneity of single unit modulation during executed and cancelled movements. **A.** No-stop trials and Stop trials were randomly intermingled on each block. Monkeys were required to touch the central target, hold the position (Hold) and wait until the Go signal (Go) instructed them to reach the peripheral target (red circle: presented either to the right or to the left; here only one position is shown). In stop trials, after the Go signal, the Stop signal (central target reappearance) instructed them to stay still (correct-stop). Wrong-stop: trials with failed inhibition. Examples of units recorded: spike density functions (SDFs) and raster plots for correct-stop trials (red) and latency-matched (control) no-stop trials (green) are shown for a specific Stop signal delay (SSD) separately for each movement direction. **C.** For the same units SDFs and raster plots from wrong-stop trials (black) and latency-matched no-stop trials (green) aligned to movement onset (M_on).

### Premotor units, associated to both movement execution and inhibition, are heterogeneous

Initially we selected those units whose activity were modulated before movement onset and/or after the Stop signal presentation in at least one movement direction compared to a control period preceding the Go signal presentation (see Material and Methods). We then considered for further analysis only those units with a stop-related modulation for at least one movement direction. Specifically, we searched for units differently modulated in the SSRT interval, that is, firing differently in the latency-matched no-stop trials and correct-stop trials, for the same movement direction (see Material and Methods). Latency-matched no-stop trials and correct-stop trials are characterized by the same maturation level of movement planning before the Stop-signal, and their comparison allows one to evaluate what happens when the movement generation is suppressed. Because of their different modulation these units can be considered predicting movement inhibition or generation.

Overall, 139 units were selected, i.e., about two third of the dataset (93 out of 113 for monkey P and 46 out of 91 for monkey C). The selected units are heterogeneous because they showed a variety of activity profiles in the two directions of movement during the task (for no-stop trials) and in relation to movement inhibition (either increasing or decreasing - after the Stop-signal - their activity in correct-stop trials when compared to latency-matched trials; see Fig. 1B) as previously described in other experiments in PMd (8, 13). Moreover, we found that about 44% (61/139) of the units presented a different profile across the two directions of movement (see Fig. 1B), while the others showed a similar modulation profile in the two movement directions. However, among the cells showing the same pattern, 49/78 (62%) was directionally selective (Wilcoxon rank-sum test, *P* < 0.05), i.e., showed a different firing rate before movement onset for the two directions of movement. Overall, directional neurons were 110 (61 + 49; 79.1%) over 139 neurons. As expected, neuronal modulation was not different in latency-matched no-stop trials and wrong-stop trials (Fig. 1 C; Fig. S2 and Supplementary Material and Methods). Overall, these results confirm previous evidence, i.e., that single units in PMd signal in heterogeneous and complex ways the action generation and suppression (13, 14).

### Population neuronal dynamics during movement execution and inhibition

The heterogeneity of the neuronal patterns observed is a confounding element in determining which aspect of the neuronal dynamics underlies movement inhibition and generation. To solve this issue, we analyzed data at the population level and reduced the dimensionality of the state space resorting to the principal components analysis (see Material and Methods for details).

Figure 2A shows, aligned to the Go signal, the average neuronal trajectories both in correct-stop (red) and in latency-matched no-stop (control trials; green) trials in the low-dimensional subspace determined by the first three principal components (PCs; explaining more than 60% of the variance in all conditions; Embedding Dimensions < 3; see also Supplementary Material and Methods and Fig. S3) for one direction separately for each monkey. As expected, the two neuronal trajectories develop similarly until the appearance of the Stop signal (empty red dot). In control no-stop trials (green lines), population activity continues describing a volley until the movement occurs (green diamond; movement onset). Conversely, after the Stop signal, the trajectory in correct-stop trials (red lines) diverges from the green one moving backward to approach the initial state. Importantly, the divergence between the two trajectories occurs well before the end of the behavioral estimate of SSRT (filled red dot).

**Figure 2.**
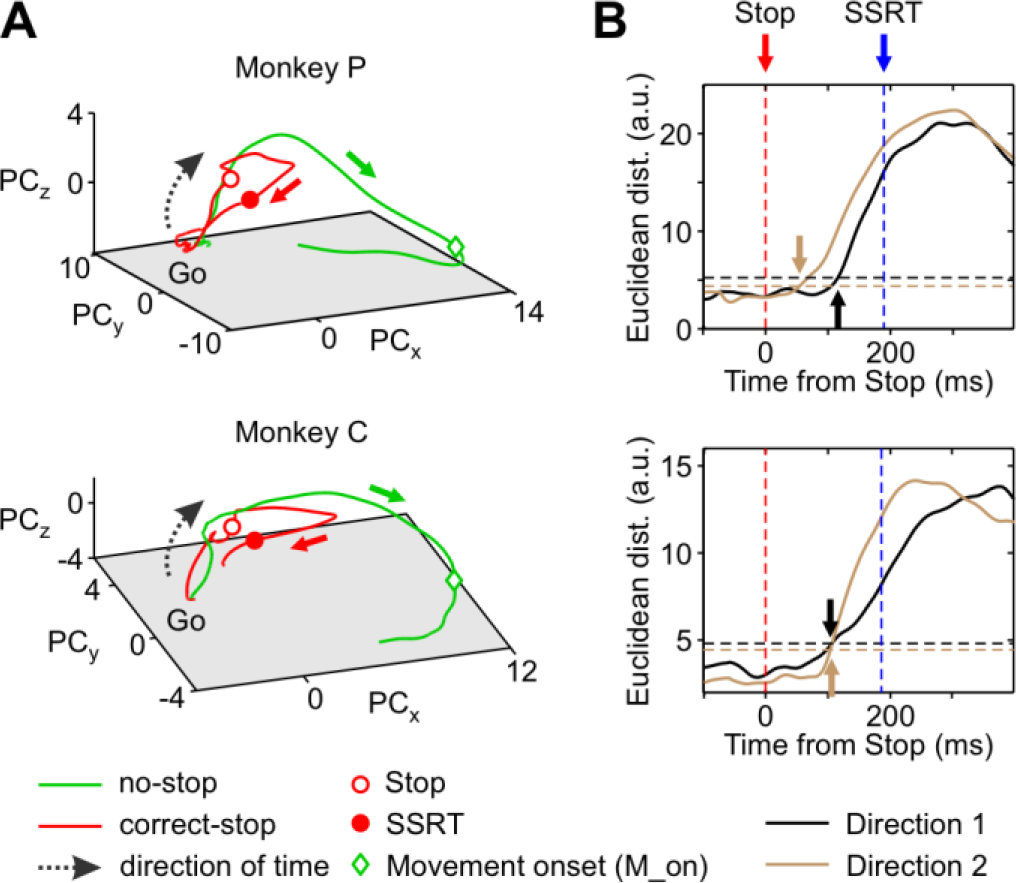
Neuronal dynamics and population estimate of the time when movements are inhibited. **A**. Neuronal trajectories of latency-matched no-stop and correct-stop trials for a single movement direction (direction 1) in the state space defined by the first three principal components. **B**. For each monkey, the average Euclidean distance between trajectories corresponding to correct-stop and latency-matched no-stop trials, aligned to the Stop signal, are represented. Arrows point out the time of significant divergence between trajectories. In Fig. 8 we refer to this divergence time as an estimate at the population level of the Stop signal neuronal time (SSNT). Horizontal dotted lines represent the reference thresholds for each movement direction (see Materials and Methods).

At a first glance, the trajectories dynamics suggests that in no-stop trials the neuronal population activity moves toward a functional state corresponding to movement generation and that movement inhibition is the active avoidance of such a process. We obtained a population estimate of the time of divergence (Fig. 2B) by calculating the Euclidean distances between the corresponding neuronal trajectories within a time window aligned to the Stop signal (from 100 ms before to 200 ms after, i.e. up to the SSRT duration; see Table S1; see Materials and Methods for further details). We found that such divergence occurred between 70 and 116 ms following the Stop signal presentation for the two directions (Fig. 2B) anticipating in time both the equivalent neuronal correlate derived from the single unit analysis (see Supplementary Results and Fig. S4) and the differences detected at the electromyographic level, when evident (see Supplementary Results and Fig. S5).

### Movement inhibition and stillness require neuronal activities to be limited into a functional subspace

We wanted to investigate whether the dynamics we observed for no-stop and correct-stop trials could be better accounted for by the existence of distinct functional subspaces. Figure 3A shows, for one animal, the average neuronal trajectories in correct-stop (red), control latency-matched no-stop (green), and wrong-stop (black) trials obtained from the first three PCs.

**Figure 3.**
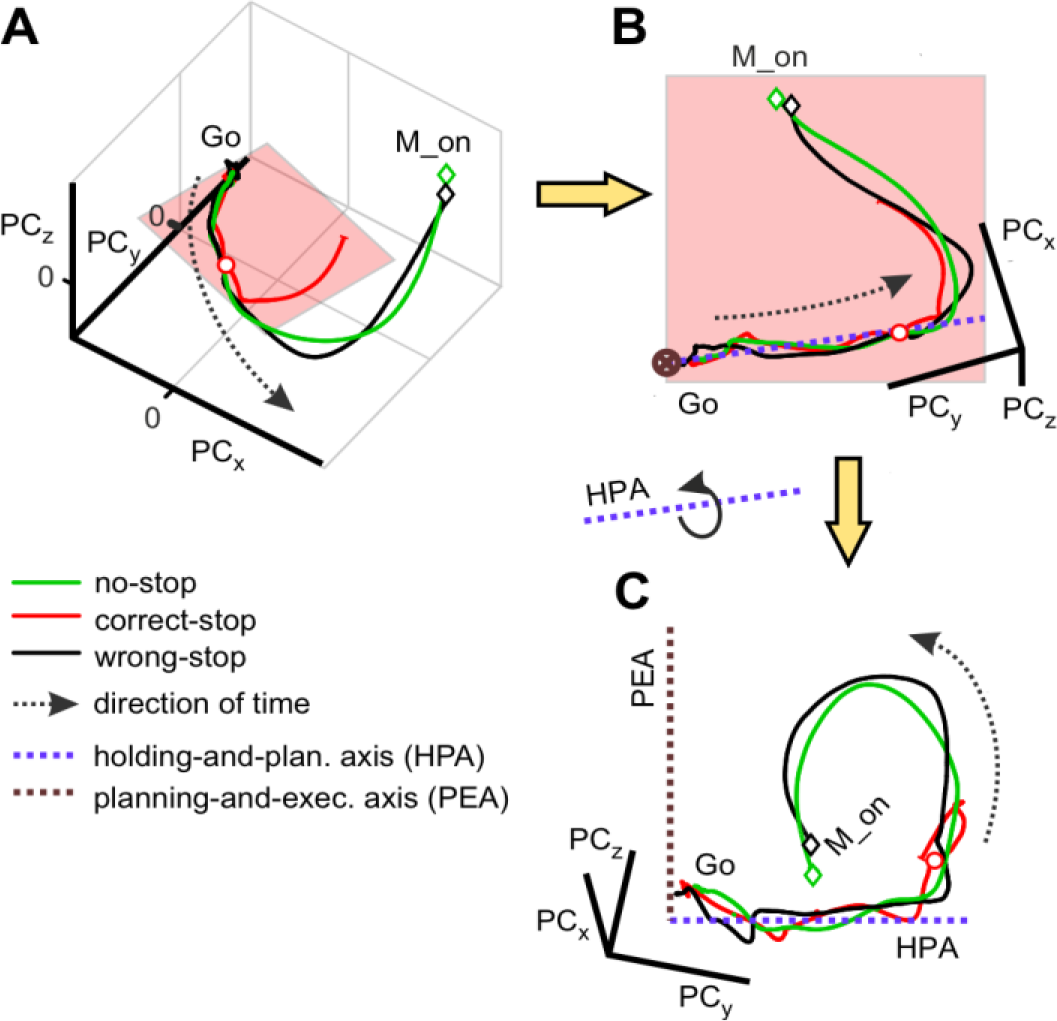
Schematic of the neuronal trajectories unfolding within the holding plane and a plane orthogonal to it. **A.** Average neuronal trajectories for all trial types aligned to Go onset. **B.** Top. Holding plane, where correct-stop trajectories mostly, reside highlighted by a rotation of the trajectories in A. The plum dotted line represents the axis which best fitted the trajectories of all groups of trials projected on the holding plane in the first 300ms following the Go-signal (hold-and-planning axis, HPA). **C**. A further rotation around the blue axis allows finding the planning and execution axis (PEA, brown dotted line).

After the Go signal, the population activity in correct-stop-trials initially follows a trajectory almost indistinguishable from the ones measured in no-stop and wrong-stop trials. The late divergence of the correct-stop trajectory (as in Fig. 2) could indicate the existence of a subspace in the neuronal state space where activities are confined before movement generation and during movement suppression. Such a hypothesis would support the generalization of the computational strategy relying on the ‘output-null’ subspace described for PMd neurons in animals tested in a delayed reaching task (6, 7).

To verify the existence of such subspace, we focused on the collective states visited by the correct-stop trials. On this data, averaged and grouped by Stop signal delays (SSD) for each animal and direction of (potential) movement. Relaying on the singular value decomposition (SVD) analysis (see Material and Methods for details), we found all trajectories of the correct-stop trials to fluctuate around a reference planar section. Figure 3B shows the view on this planar section of the average trajectories in Fig. 3A here obtained, for illustrative purposes, after a suited rotation of the original 3D space. In this plane the similarity among no-stop, wrong-stop, and correct-stop trajectories is remarkable. Thus, the dimensionally-reduced population dynamics highlighted from this perspective does not contain any information about whether or not the movement will be generated.

However, as premotor cortices are known to also represent the motor output in reaching tasks (2), a related activity subspace must exist to allow independently and robustly reading out these motor commands from the activity evolution of the network (7), and thus allowing to distinguish between trials where movement is executed or inhibited. To this end, we extracted from the same plane an axis by further fitting data from correct-stop trajectories in the 300 ms following the Go-signal (Fig. 3B). This axis was then used to obtain a plane orthogonal to the first one. Figure 3C shows (again for illustrative purposes only) this orthogonal plane, together with the corresponding dynamics of all average neuronal trajectories. From this last perspective, it is rather apparent that movement related activities (no-stop and wrong-stop trajectories) depart from a reference holding plane where correct-stop activities are instead confined for most of the time.

We called the first axis the ***holding-and-planning axis (HPA)***, and the second subspace the ***planning-and-execution axis (PEA).*** The reason for these definitions will be apparent in the following.

### Movement generation as an escape from movement inhibition subspace

Our first step of analysis suggested the presence of two subspaces, where decision-related neuronal dynamics unfolds differently. Thus, some questions arise: how can these segregated dynamics explain movement generations and suppression? What makes a stop trial correct or wrong? To address these issues, we inspect in more detail how the projections of the neuronal trajectories onto the described axes change in time.

Figure 4A shows (grouping trials by either RT or SSD for wrong/no-stop trials and correct-stop trials, respectively; see also Fig. S6) that neuronal activities, observed as projections in the PEA, immediately after the Go signal are roughly stable, slightly fluctuating around the reference zero-baseline, and then they move towards more negative values, ‘drawing’ a trough. During the same period, neuronal activities observed as projections in the HPA (Fig. 4B) show a peaking ramp-like dynamics, similar for all trials and conditions (see also Fig. S6).

**Figure 4.**
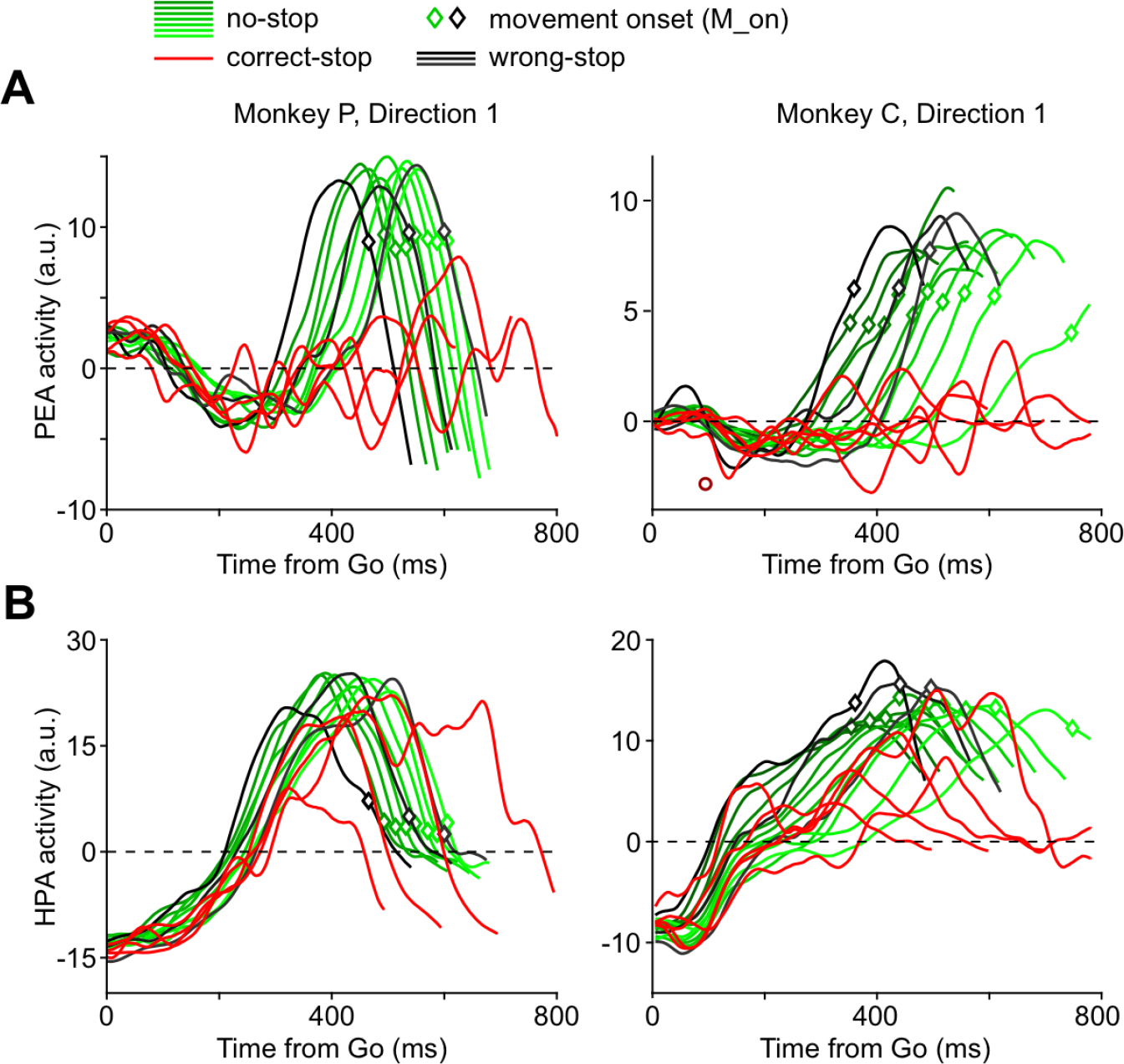
Components of the neuronal dynamics determining RTs and correct-stop trials. A. Projections on the planning-and-execution Axis (PEA) of the neuronal trajectories (direction1) – namely, PEA activity – grouped by RTs (no-stop in deciles and wrong-stop trials in tertiles) or SSDs (correct-stop trials). After an initial period around the Go signal in which trajectories are overlapped, no-stop and wrong-stop trials projections diverge from correct-stop trials projections. B. Projections on the holding-and-planning Axis (HPA) of the same neuronal trajectories – namely, HPA activity – as above: in this case the projections display an initial ramp-like dynamics and similar features for all trials and conditions.

In a phase successive to the troughs, Fig. 4A shows that PEA activities, in particular those related to wrong-stop (black lines) and no-stop trials (green lines), escape from the plane signaling an important change in the overall state. In the HPA, as described above, it is instead very difficult to differentiate among the different behavioral outcomes.

Projections on both HPA and PEA show a strong correlation with RTs (see Fig. S7). The relationship was slightly stronger when considering projections on PEA in ¾ of the conditions [median beta coefficient (interquartile range, IQR): monkey P, direction 1: PEA = 0.99 (0.03); HPA = 0.91 (0.38), p < 10^-4^; direction 2: PEA= 0.98 (0.07); HPA = 0.89 (0.43), p < 10^-4^; monkey C, direction 1: PEA = 0.99 (0.02); HPA = 0.81 (0.40), p < 10^-4^; direction 2: PEA= 0.90 (0.07); HPA = 0.95 (0.62); p = 0.1]. The higher IQRs suggest a more variable relationship of projections onto the HPA with RTs.

We further asked in which of the two projections the relationship with RTs first emerged. Interestingly, we found no evidence of correlation with RT during the post-Go initial phase of neuronal modulation in the PEA (i.e., during the descending phase, toward more negative values). In fact, we found that in HPA the significant relationship between neuronal activities and RTs emerged earlier than in PEA [mean (SD); monkey P, direction 1: HPA = 377 (246) ms, PEA = 482 (133) ms; direction 2: HPA = 300 (278) ms, PEA = 450 (123) ms; monkey C, direction 1: HPA = 276 (104) ms, PEA = 332 (101) ms; direction 2: HPA = 299 (182) ms, PEA = 365 (72) ms; rank-sum test p < 10^-4^ in all cases]. Thus, although in both projections a significant relationship with RTs emerged, this relationship developed as a chain first involving the HPA projections and only afterwards involving the PEA projections.

### Ramps in HPA contribute to motor planning without relying on a threshold mechanism

The ramp-like dynamics after the Go signal in the HPA (Fig. 4B) is typically observed in decision processes or when an internal generated timing signal is employed to elaborate an input signal (15, 16) and is associated with a threshold mechanism. In such threshold mechanisms the amount of activity, *per se*, is sufficient to determine the transition towards a different state once a certain level is reached. Concurrently, a pre-movement reduction in neuronal variability is usually observed, deemed to correspond to the termination of a decision-related process (2, 3).

Figure 5A shows, on one hand, that when activities escape from the reference plane in PEA and a movement is generated (wrong-stop trials and no-stop trials; empty black and green squares respectively) the activity in HPA tends to stay close to a subspace where neuronal variability shrinks (see also Fig. S9). On the other hand, it is evident that one cannot predict movement onset relying only on this level of activity in the HPA. In fact, activity of correct-stop trials (red traces and empty red circles) can be even higher than the activity observed in no-stop trials in the HPA. Conversely, Fig. 5B shows the presence of a threshold when activities are observed in the PEA. Indeed, when movements are inhibited, the projection of activity in the PEA (red traces) tends to remain under the level necessary for muscle activation. This level is instead reached by both no-stop and wrong-stop trials activities (green and black dots, respectively).

**Figure 5.**
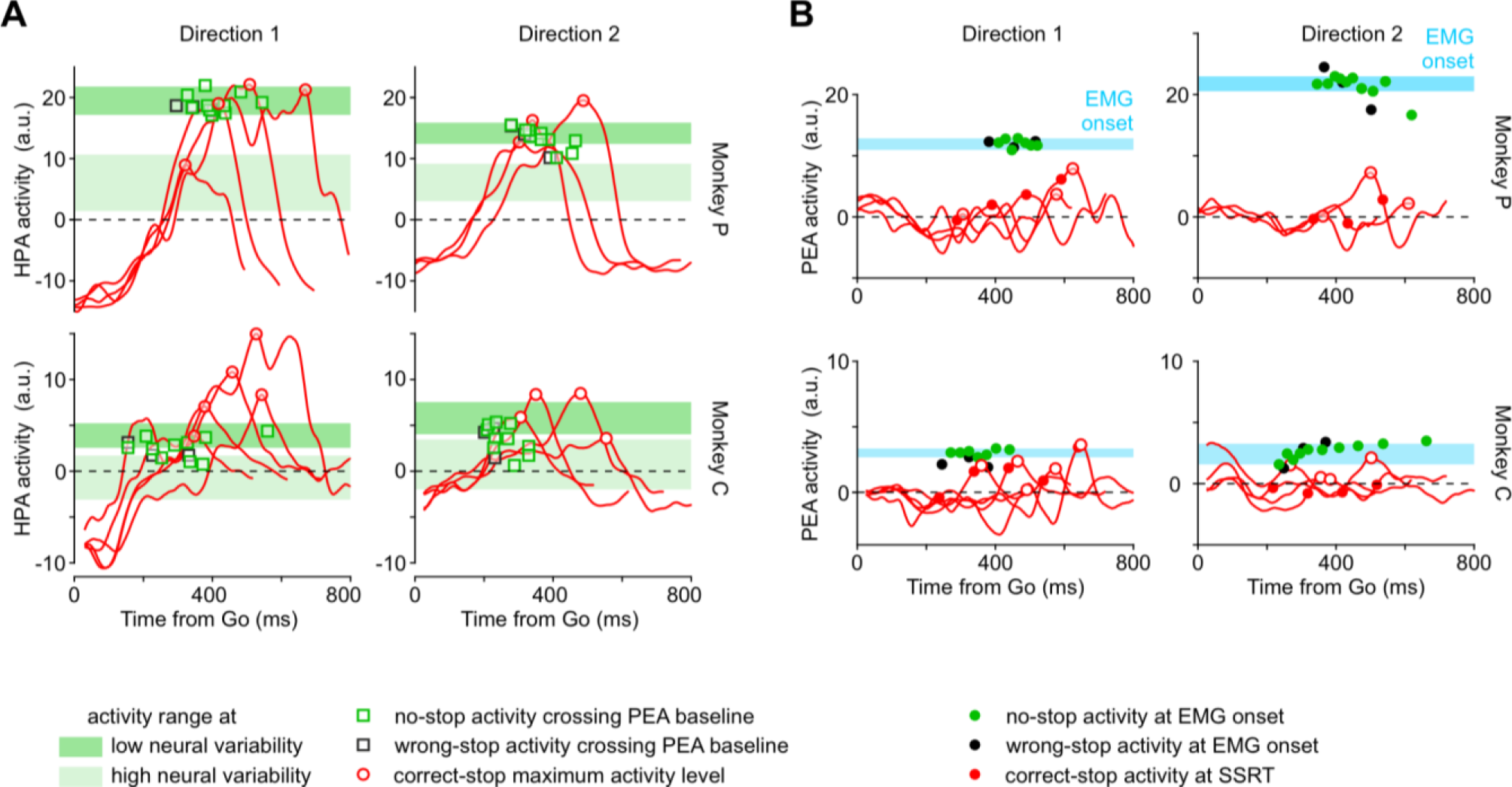
Dynamics for movement generation exploits a threshold-based mechanism in the PEA only. **A**. Projections of correct-stop activities in the HPA as in Fig. 4 (red traces); the maximum activity level is indicated by the empty red circle. Green and black squares indicate the HPA activities for no-stop and wrong-stop, respectively, measured at the times when projections on PEA of the same activities crossed the zero baseline (see panel B). The shadow bars represent the levels of no-stop activity (mean ± SD) measured in the HPA when the neural variability across trials (Fano factor) was high (light green) or low (dark green) before movement onset (see Fig. S9). B. Projections of correct-stop activities in the PEA as in Fig. 4 (red traces); the maximum activity level is indicated by the empty red circle. Filled red circles correspond to PEA activity of correct-stop trials at SSRT. Green (black) filled circles display the PEA activity during no- stop (wrong-stop) trials at the time of EMG onset. Cyan bars represent the range of activities projected in the PEA when muscles are activated before movement generation (mean ± SD; calculated on the muscle’s latency confidence interval, CI).

Why does the ramp-like increase of activity in the HPA, as shown in the previous paragraph, resulted highly correlated to the RT? We suggest that this ramping activity is needed to bring the system into a state necessary (but not sufficient) to elicit the motor-related activity in the PEA, thus encoding an essential part of the movement plan. To corroborate such a hypothesis, we used a different paradigm in monkey P (see Fig. S11). There the animal was informed whether the movement was possible, certain, or not required, by presenting a cue well before the presentation of a Go or NoGo signal. We found that activity in HPA only ramps when movements are possible or certain. Indeed, when the movement is not required, even the presentation of the Go signal does not induce any significant change in the HPA. Thus, the dynamics we described in HPA cannot be considered a mere lack of movement related activity but, instead, an active process involved in both initial position holding and planning of the incoming movement.

### Movements are generated when PEA activities cross the boundary of the holding subspace

As can be expected from Fig. 4, PEA activities for no-stop and wrong-stop trials are also highly stereotyped in time when aligned to the movement onset (Fig. 6). It is important to remark here that this dynamics occurs while specification of movement direction is already determined (see Fig. S10).

**Figure 6.**
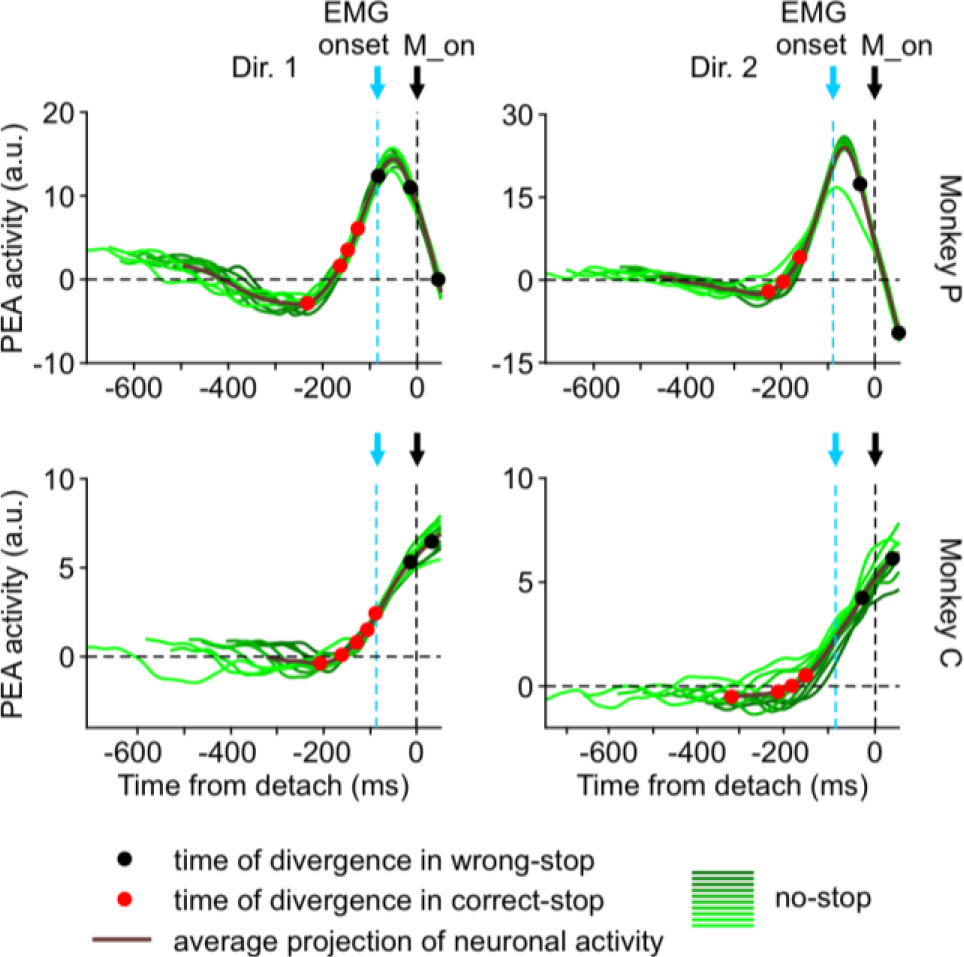
Movement inhibition is effective only if the Stop signal is processed before full muscle contraction. Each panel represents the PEA activity for the no-stop trials (thick lines: average; green lines deciles) aligned to the movement onset. In the average activity, dots highlight the activity at the time of neuronal divergence for correct-stop (red) and wrong-stop (black) trials. The timing of the neuronal divergence is computed as in Fig. 2B relying on single values of SSD for correct-stop and on the average SSD for wrong-stop trials.

Conversely, the PEA activity during correct-stop trials (Fig. 4A) does not deviate too much from the baseline. This is compatible with the hypothesis that correct-stop activity trajectories are trapped into a specific subspace as in these trials movement planning is not yet mature at all or it is cancelled well before its complete maturation, compatible with what preliminary observed in (4, 9).

In this framework, a successful Stop trial could be obtained by temporarily suppressing the translation of the motor plan into an overt movement, i.e., avoiding the escape from the trough in the PEA. Thereafter, movement onset is possible only if activities also cross a point-of-no-return, represented by the muscle activation. This picture is confirmed in our experiments as the Stop signal can be effectively processed only if occurring before muscle contraction (Fig. 6). This is, once again, clearly reminiscent of a threshold-like mechanism. Indeed, the PEA activity is higher when a movement is performed (Fig. 6, black circles before movement onset), compared to the activity at the time of neuronal divergence in correct-stop trials (Fig. 6, red circles).

Note that PEA activity in Fig. 6 was confined in the trough for variable durations before the movement onset. In Fig. 7 we quantified the relationship between such time intervals and the RT finding that the exit times (Fig. 7A), are highly predictive of the movement onset (Fig. 7B; see also Fig. S11B). Movement generation is then kept at bay for so long as neuronal activity is confined into the trough below the reference subspace (negative values observed: mean ± SD, Monkey C: Dir.1= - 0.5 ± 0.3; Dir.2 = -0.59 ± 0.36; Monkey P: Dir.1 = -2.0 ± 1.0; Dir.2= -1.6 ± 1.1, t-test p < 0.01 for all cases).

**Figure 7.**
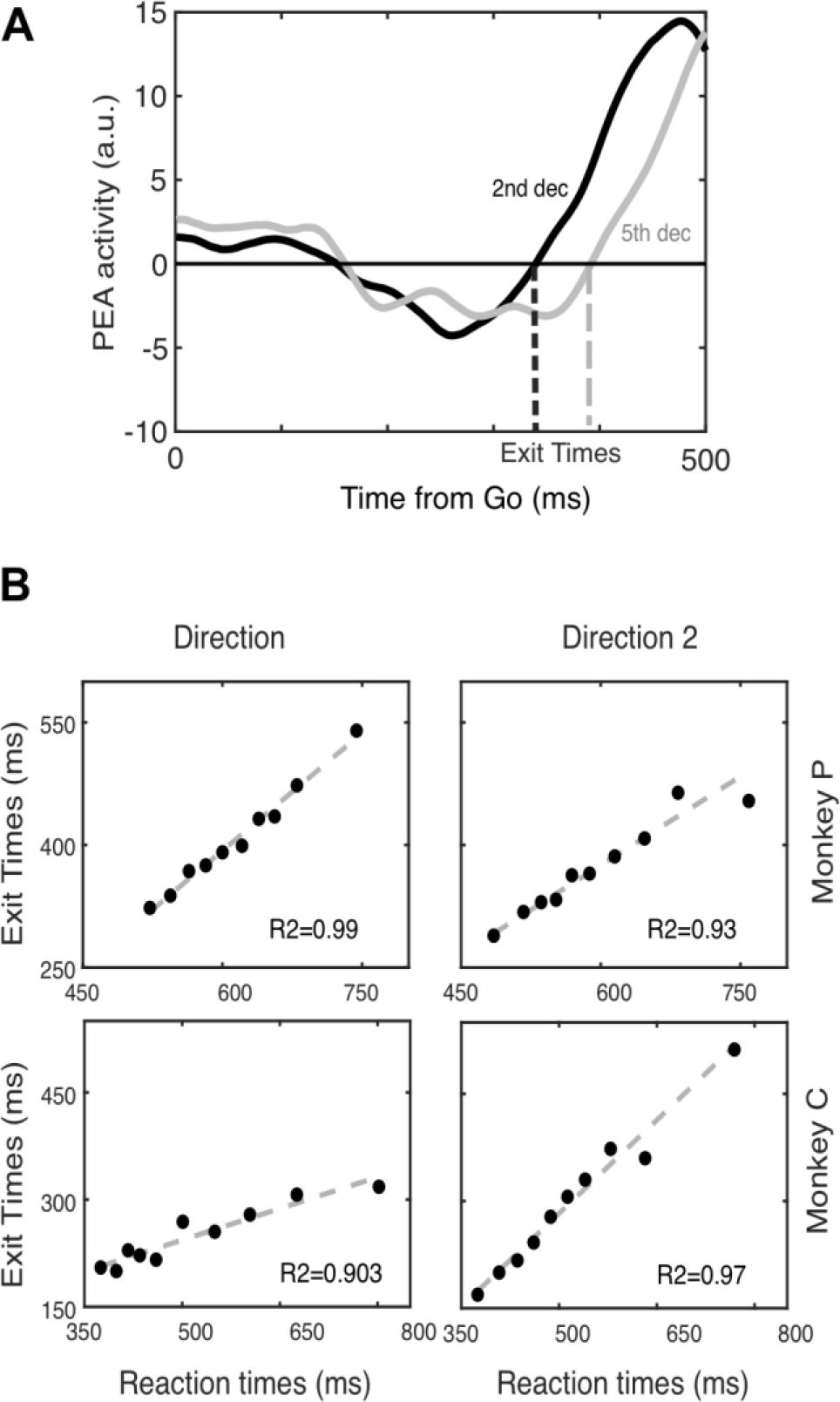
Exit Time estimate and relationship with reaction time. **A.** Schematic of the exit time computation for two deciles in the dataset. **B.** Correlation between Exit Time occurrences and RTs for each movement direction, separately for each monkey.

Overall, the emerging picture is that after target onset the neuronal dynamics visits different states better differentiated when observed as distances from different subspaces. The movement is generated when this activity grows reaching a threshold (see Fig. 5B). Importantly, this growth of activity in the PEA starts only when the activity reaches a certain level in HPA, purportedly corresponding to the end of a decision process. However, the reach of this level in HPA is not sufficient to determine the movement onset. Indeed, it is reached also in correct-stop trials when there is no movement production.

## Discussion

In the present study we investigated the premotor neuronal dynamics underlying the control of arm movements by combining two complementary lines of research: i) neuronal population analysis, by exploiting a state-space approach (see for example 6, 17, 18), and ii) the countermanding task, an experimental design ideally suited to investigate and model volitional control of movement (19, 20).

First, we confirmed that PMd units modulate their activity before the end of the SSRT thus having temporal characteristics that allow predicting whether a movement will be halted (8). Secondly, we further extended this finding by showing that neuronal modulation precedes muscle modulation in successfully inhibited trials, strongly supporting a causal role for this area in movement generation. Thirdly, and most importantly, by applying dimensionality reduction techniques, we uncovered the existence at the population level of a new holding subspace, which we suggest generalizing the previously introduced ‘output-null’ subspace (6). In our view, this subspace is where premotor neuronal activity must actively be confined to avoid the transition of motor plan into execution. Such novel dynamics behind the control strategy extends the concept of the ‘output-null’ subspace, and it is complementary to the one adopted in previous works on delayed reaching tasks.

### Movement control in a state-space framework

The main goal of this study was to uncover the premotor neuronal dynamics that characterizes movement generation by contrasting neuronal activity during movement production with neuronal activity during active movement inhibition. We found that following the Go signal the neuronal activity is confined into a (holding) subspace where it is initially similar between correct-stop, wrong- stop and no-stop trials. In this initial stage the neuronal activity is ‘trapped’ into a trough, visible from a specific perspective, possibly reflecting the specific task context that is required to block movement execution (see also the control task results; Fig. S11). As the neuronal dynamics evolves a clear differentiation emerges: in no-stop and wrong-stop trials the neuronal activity moves away from the trough towards a state-space region which almost deterministically anticipates the movement generation. Differently, in correct-stop trials (where no movement is produced) the neuronal activity remains confined in the initial subspace, separate from the trajectory observed in no-stop trials. Importantly, the temporal evolution in no-stop trials (and wrong-stop trials) is strongly related to RTs, and it shows a clear stereotyped nature when aligned to the movement onset. These pieces of evidence demonstrate the existence of neuronal dynamics that anticipates motor behavior clearly distinguishing the active inhibition versus the movement generation. Thus, we conclude that PMd expresses specific neuronal dynamics underpinning either movement generation or inhibition. This conclusion is further strengthened by the fact that the divergence between no-stop and correct-stop trials occurs before the end of the SSRT (21).

Our data are in line with a series of findings obtained from studies investigating the preparation and execution of reaching movements by using state-space approaches (6, 22–24). In most of these experiments monkeys are provided with prior information about movement direction by a cue followed by a Go (move) signal after a delay. During the preparation phase (delay period) a cascade of neural events bring the collective activity into a prepare-and-hold (attractor) state (2, 4, 17). As proposed, the goal of this state is to set the motor plan, i.e., a neuronal state which functions as the initial condition determining the upcoming movement (2, 24, 25). The neuronal trajectories will then evolve towards a complementary subspace, but linked, to the previous one (7). In this last subspace, reaching execution dynamics (rotational) unfolds (18). Within this framework the preparatory activity does not generate the movement *per se* but it always occurs before the movement onset, even when very brief times to prepare the movement (zero delay) are available (6, 23, 26). The transition from preparation to execution is characterized by some degree of overlapping between preparation and movement phases (23), and by a strong condition-invariant signal that precedes the movement onset (27).

In summary, an important aspect of the previous studies is that the generation of the movement was always required, thus it is not clear which aspect of the dynamics can be deemed necessary for movement generation. We employed the countermanding task to address this issue. To this purpose we exploited the concept of neural manifolds (28) finding a holding subspace where neuronal trajectories of the correct-stop trials, as well in the initial part of no-stop and wrong-stop trials, are confined. The escape of neuronal trajectories from this subspace determines the transition from the prepare-and-hold state to the movement generation. This dynamics is highly stereotyped and precedes the movement onset by a fixed time-lag, similar to what is observed in other studies (27). This dynamics could correspond to the information to start the movement, an internal Go signal, that PMd can provide to other cortical and subcortical structures, as previously suggested (6, 7).

This allows to avoid searching for specific inhibitory circuits that, so far, have been impossible to define in the premotor and motor cortices of primates, due to the difficulties in accounting for the connections of cortical neurons to subcortical, brainstem and spinal cord neurons (29), although recent attempts have been promising (30).

### Nonlinear network dynamics underpinning the holding subspace

HPA and PEA are orthogonal axes, meaning that activity projection on such axes is mainly due to different subsets of units (Fig. 1B). In other words, ramping activity along the HPA is probably due to a subset of units displaying a post-Go quasi-linear increase/decrease of the firing rate (sketched on Fig. 8A-bottom). Instead, the sudden increase of PEA activity tightly locked to the movement onset is determined by a pool of units displaying sharp transitions from low-to-high activity levels and *viceversa* (see Fig. 8A-top).

**Figure 8.**
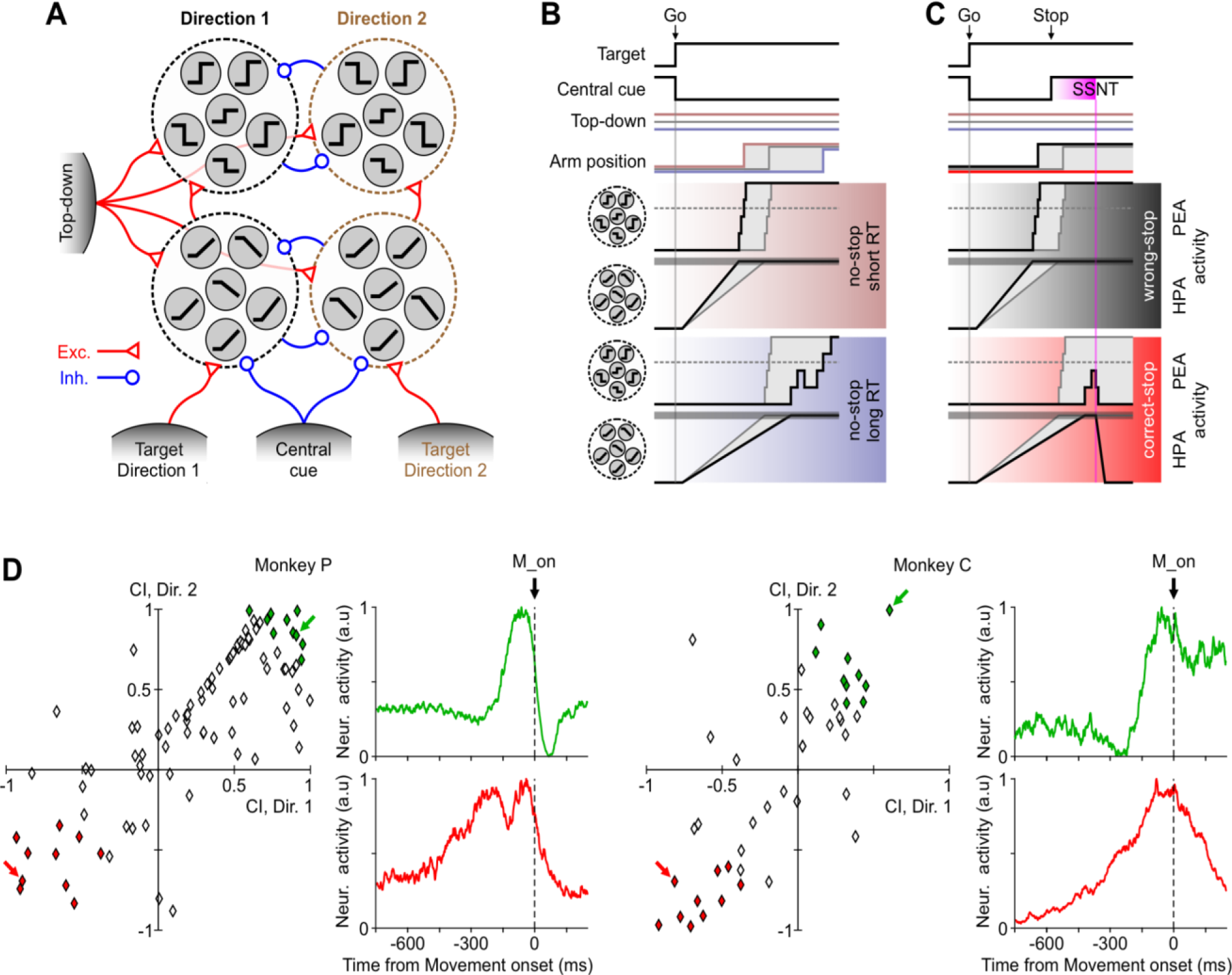
Schematic proposal of PMd organization underlying movement execution and inhibition. **A.** Pools of ramp-like and switch-like units (grey circles) respond differently depending on the target to be reached, signaled by a selective excitatory input coming from other brain areas (grey shaded units at the bottom). Target related representations are mutually exclusive due to cross inhibition. Ramp-like units contributing to the rising of HPA activity elicit sharp- transitions in switch-like units. The latter contributes to the changes of PEA activity that indirectly react to the target-related input. A unspecific top-down input (left) making differently excites all the PMd units from trial-to-trial. Onset and offset of the central cue signaling to Stop and Start, respectively has an inhibitory effect on the PMd units. Red and blue arrows represent excitatory and inhibitory connections, respectively. **B.** Behavioral, environmental (top) and neuronal changes in no-stop trials with long (bottom) and short (middle) RTs, which are determined by a relatively weak (bluish) and strong (reddish) top-down excitation, respectively. Grey traces represent a reference trial with average RT. Black traces, pooled activity of switch-like and ramp-like units are plotted in black side by side with the respective unit icons. These traces are those resulting from projecting the high-dimensional population activity on the PEA and HPA, respectively. Horizontal dashed grey lines, activity level when movement onset is irreversibly elicited when crossed in the PEA. Simultaneously, when in the HPA the pooled ramp-like units reach the horizontal gray strip, the switch units are facilitated to change state. **C.** As in panel B, for those trials in which the Stop signal is presented. Black and red shaded subpanels correspond to example wrong- and correct-stop trials, respectively. SSNT, stop-signal neuronal time (corresponding to the time of significant divergence between trajectories after the Stop-signal, as in Fig. 2B). **D**. Scatterplot of contrast indexes (CI) for each movement direction in both monkeys. CI = (HPA_weight_ − PEA_weight_)/(HPA_weight_+ PEA_weight_) where the “weight” is the element associated with a single unit in the corresponding axis (see Materials and Methods). Green and red diamonds represent units with extreme values in PEA and HPA contributions respectively. Example units of the extreme groups, highlighted by colored arrows, are shown using the same color code of the scatterplots.

In the specific framework investigated here, the relatively slow and linear drifting along the HPA determines a post-Go ‘refractory phase’ where presumably target-related information accumulates, and no movement occurs (Fig. 8B and C). The passage across a region (see Fig. 5A, low neuronal variability) after a variable time elicits the rise of the PEA activity. In those trials with short RTs (Fig. 8B and C - middle), the related sharp transitions of that activity are strongly stereotyped even in the Go-centered profiles (Fig. 6), meaning that units displaying a fast switching of activity are rather synchronized in time (see also 4).

In correct-stop trials (Fig. 8C, bottom) and in no-stop trials with long RTs (Fig. 8B-bottom) more gradual HPA ramps and less stereotyped PEA activations of the switching units are visible. Intriguingly the latter appears to fluctuate up and down (Fig. 4A), and under this condition, motor program maturation can be then interrupted if it is not fully developed, i.e., the activity is confined below a trigger threshold (Fig. 8C-bottom). Indeed, only when planning has reached a sufficient degree of maturation it can be translated into an overt movement.

To test the existence of two segregated sets of neurons (Fig. 8A), we compared the contribution to the PEA and to the HPA of each single unit, finding a continuum of neuronal types (Fig. 8D) which is preserved across movement directions. However, a subset of these neurons displayed a rather polarized role showing either highly positive or negative contrast indexes. In our model, they represent the units having sharp (see green units in Fig. 8D) or ramping-like activity (red units), respectively. These two neuronal types appear to be universal components in cortical areas involved in arm movement planning and motor decision (4, 15, 16, 31).

RTs are both determined by the time needed to reach an accumulation threshold (dashed line in Fig. 8B) and by a variable time in which switching units complete the transformation of the cortical state into the one encoding the motor plan. The latter phase has a more prominent role only for relatively long RTs, and the Stop signal can be successfully processed both if SSRT falls after the end of the first accumulation stage. In possible short-RT trials the onset of Stop signal must be early in correct-stop trials such that the SSRT could precede the beginning of the activity rise along the PEA. If one of these two conditions is not fulfilled, a wrong-stop trial is performed (Fig. 8C-middle).

A mechanistic implementation of the discussed interplay between ramping and switching units on one hand requires that the former provide input to the latter, such that once the input received by the switching units crosses a trigger threshold, a cascade of activity switches occurs (4, 32). On the other hand, one should also explain the trial-by-trial variability of such chain of reactions, which underlies both the capability to inhibit an instructed movement and the variability of the RTs. To this aim, an additional unspecific input has to be taken into account. Indeed, a tonic input modulating the excitability of both unit types would allow changing from trial to trial the switching rate and the ramp slope (33, 34). This input could be provided by a proactive tonic top-down control, which varies across trials (top-down signal in Fig. 8A). Finally, we remark that ramping activity in principle can be obtained by pooling time shifted switch units, just to mention the possibility to have the latter kind of neurons contributing to both pools sketched in Fig. 8A (35, 36).

The existence of such a proactive control mechanism in models of saccadic countermanding tasks is not new (37, 38). For instance, in (37) a top-down control of frontal origin implemented a post-Go phasic reduction of the excitatory input to a specific subset of neurons. This sudden change in the input resembled the effects of a homunculus intervening at specific random times aiming at recovering the RTs statistics and inhibition function (19). To work around such limitations, it has been suggested that another proactive mechanism is at work leading to a modulation of the ‘response caution’ governed by the baseline activity of the inferior-frontal gyrus (IFG) (38). This in turn, slows down or accelerates the movement production via the IFG-subthalamic nucleus hyperdirect pathway. Our results are compatible with this second scenario, as each trial has its own degree of excitability/stability of the ramping and switching units that we suggest being associated with a different top-down tonic input (i.e., the IFG baseline activity). Differential stability of the no-movement condition can be interpreted as a different speed of the Go process in the race model (39, 40), and we predict it is related to the high-beta state found in (41) associated to the capability of the cortical-basal ganglia circuits to stabilize selected motor plans.

Under this hypothesis, the permanence in the newly found holding subspace (i.e., represented by the trough in PEA activity) appears to be effectively separated from the boundaries delimiting the previously introduced output-null space (6). This in our view is obtained by making the holding subspace a more “attracting” region. In the framework we propose, the attracting force is modulated by an unspecific top-down control, while the premotor network can autonomously implement the machinery needed to produce and inhibit an instructed movement. In turn, the stability of this holding state allows having complex and computationally relevant population dynamics such as the accumulation process preceding the full maturation of the motor plan, without the danger to cross the boundary of the output-null subspace triggering the movement onset.

## Material and Methods

### Subjects

Two adult male rhesus macaque monkeys (Macaca mulatta; P, C) weighing 7–9.5 kg served as subjects. All experimental procedures, animal care, housing, and surgical procedures conformed with European (Directive 86/609/ECC and 2010/63/UE) and Italian (D.L. 116/92 and D.L. 26/2014) laws on the use of nonhuman primates in scientific research and were approved by the Italian Ministry of Health. In both monkeys a 96 channels Utah arrays (BlackRock Microsystem, USA) was implanted using anatomical landmarks (arcuate sulcus - AS - and pre-central dimple - pCD) after dura aperture on the dorsal premotor cortex (PMd) contralateral to the arm employed during the experiments (Fig. S1E)

### Apparatus

Stimuli were presented on a 17-inch LCD monitor (800×600 resolution) equipped with a touch-screen (MicroTouch, USA). Stimuli consisted of red circles with a diameter of 2.8 cm on a dark background. During the performance of the task eye movements were monitored by using a non-invasive Eye-tracker (Arrington Research Inc, AZ).

### Behavioral task

Monkeys performed a countermanding task (Fig 1A). They had to touch the stimulus with their fingers, hold and fixate it (Holding Time range: 400-900 ms). Thereafter the central stimulus disappeared and, simultaneously, a target appeared (Go signal) randomly at one of two opposite peripheral positions (left and right) instructing for a movement in the corresponding direction (indicated as direction 1 and direction 2). In no-stop trials to get a juice reward, monkeys had to start the arm movement within a maximum time (1000 ms) to discourage from adopting a procrastination strategy due to the presence of stop trials, and to maintain their fingers on it for a random time (400-800 ms, 100ms step). In stop trials at a variable delay (Stop signal delay, SSD) after the go signal was presented, the central stimulus reappeared (Stop signal) instructing the monkey to keep the hand on the starting position (additional holding time; 400-1000 ms interval, 100ms step) to perform a correct-stop trial and earn the juice. If the monkey moved the hand during stop trials, the trial was considered a wrong-stop trial and no reward was given. No-stop and stop trials were randomly intermingled in such a way that the no-stop trials were more frequent (66%).

Data were collected using a staircase tracking procedure to change the SSD from one stop trial to the next according to the behavioral performance: if the monkey succeeded in withholding the response (correct-stop trial), the SSD increased by a fixed amount (100 ms); if it failed (wrong-stop trial), the SSD decreased by the same amount of time. The goal of the tracking procedure is to determine a SSD for which the probability of response (i.e. the probability to have a wrong-stop trial) is 0.5 or very close to this value. This allows estimating a reliable stop signal reaction time (SSRT; 40, 42, 43), see below for further details).

### Behavioral analysis

The countermanding task permits estimating the Stop signal reaction time (SSRT) by extracting three main variables: the reaction time (RT) distributions of no-stop trials and wrong-stop trials, and the probability to respond [p(R)] by error to the Stop signal. These data are modelled according to the race model (39) to establish first if the assumptions of the model are respected (see below) and, in this instance, to estimate the SSRT. Briefly the model states that in stop trials two stochastic processes race toward a threshold: the Go process, started by the Go signal, and the Stop process, started by the Stop signal. The behavioural result of this race, either movement generation in wrong-stop trials or movement suppression in correct-stop trials, will depend on which of these processes first will reach his own threshold. In correct-stop trials the Stop process wins over the Go process and vice versa. By changing the SSD the output of the race is affected: the longer the SSD, the higher the probability to facilitate the GO process.

An important assumption of the model is that the Go process in the stop trials is the same as in the no-stop trials (independence assumption; 39, 40, 43, 44; see Supplemental Material and Methods).

We calculated the SSRT by using the integration method because it is the most reliable (44). This method requires subtracting a given SSD from the finishing time of the STOP process. The finishing time of the STOP process is calculated by integrating the no-stop trial RT distribution from the onset of the Go signal until the integral equals the corresponding observed proportion of wrong-stop trials for the given SSD (45). We obtained the estimate first by calculating the mean SSD presented in each session and then subtracting it from the finishing time obtained by considering the overall p(R) (44, 46, 47).

### Neural recordings

The unit activity was first isolated online, and then controlled off-line by using specific software (Open Sorter, Tucker Davis Technologies). In this work we included both single unit as well multi-unit activity. Multi-unit was defined as spiking activity that could not definitively be attributed to a single unit. In the text we use the term unit without distinction. This is because the main analyses we performed are based on population activity and dimensionality reduction techniques typically produce very similar results regardless of the employment of single or multi-unit (27, 48). Array data in this paper come from a single recording session for both monkey P and monkey C. The use of data coming from a single, not repeated recordings, allowed us to completely avoid risks of unit replication across different sessions.

### Neuronal analysis at the single unit level

To perform some of the neuronal analysis, we represented the neuronal activity by a spike density function (SDF) obtained by convolving the spike train with an exponential function mimicking a postsynaptic potential: we used the same equation described in (49) where the convolution kernel K(t) of the neuronal activity is:

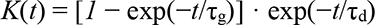

where τg = 1 ms corresponds to the growth phase of the synaptic potential, and τd = 20 ms is the decay phase (50).

We selected for analysis units that showed an increase/decrease of the average firing rate before movement onset (pre-RT, from -200 to -50ms before movement onset), compared to the 200 ms before Go signal (pre-Go) for at least one movement direction (Wilcoxon rank-sum test *P* < 0.01), and/or on units that showed a significant difference between correct-stop trials and latency-matched (see below) no-stop trials after the Stop signal. For this last case average neuronal activities in 50 ms not overlapping bins were compared in the (+100, +400) ms interval after Stop signal presentation. A neuron was included for further analysis if it showed a significant difference for at least one bin (Wilcoxon rank-sum test *P* < 0.01). These analyses were performed by using the exact spike counts. Directional selectivity was evaluated by comparing the pre-RT activity between movement directions (Wilcoxon rank-sum test *P* < 0.01).

For correct-stop trials the latency-matched no-stop trials were those with RTs longer than SSD+SSRT for each specific SSD and session; for wrong-stop trials latency-matched no-stop trials were those with RTs shorter than SSD+SSRT.

For each unit, we derived the time lag between the stop-related neuronal latency and the behavioral estimate of successful inhibition (i.e., the SSRT). The stop-related neural latency was estimated by subtracting the SDF of correct-stop trials from that of latency-matched no-stop trials considering the time when the differential SDF exceeded by 3.5 SD (or fall behind by -3.5 SD) the mean difference in baseline activity calculated from −400 ms to the time of Stop signal, and remained above (below) this threshold for at least 50 ms. A negative time lag indicated a stop-related neuronal latency that occurred before the end of the SSRT.

### Neuronal population analysis in a low-dimensional state space

To study the neuronal dynamics at the population level, we resorted to a standard principal component analysis (PCA) to reduce the dimensionality of the recorded state space. The approach we developed consists of two main steps. Firstly, we characterized the neuronal dynamics of the movement suppression process and the latency of its neuronal correlate at the population level. Then, we evaluate the planning and movement generation process across different trial types and RTs.

In the first step we calculated average spike density functions in 1-ms time bins by aligning the trials to the Go and to the Stop signals, separately. For no-stop trials where the Stop signal was absent, we considered the hypothetical SSD corresponding to the average SSD obtained for correct-stop trials. As results, for each trial type and movement direction the spike densities from all the different units composed a matrix of samples. We then concatenated these three matrices preserving the number of rows (N = number of units). The activity of each unit was then normalized by subtracting mean activity and dividing the activity standard deviation computed across all conditions. The obtained neuronal activities (matrix rows) were eventually smoothed with a Gaussian kernel of 50-ms width. In this framework, the population state of the probed cortex at any time was represented by a N-dimensional vector (a column of the aforementioned activity matrix).

At this stage, we computed the distance between average no-stop and correct-stop trials trajectories to estimate the time of divergence at population level. More specifically, in the above N-dimensional state space we carried out the Euclidean distance between no-stop and correct-stop trials at each 1-ms time bin in the interval (-100, 200) ms centered around the Stop signal. This was done for each movement direction. We then calculated the mean and the SD of these distances and we set a threshold level corresponding to mean + 3 SD. The latency of the Stop process onset was the first time from the Stop signal onset when the population activity distance exceeded the threshold level.

To capture the inter-trial variability associated to different behaviors and at the same time to reduce the unavoidable fluctuations of the estimated spike densities, we further divided the no-stop trials into ten groups (deciles) and the less numerous wrong-stop trials into three groups (tertiles) based on the ordered RTs (neuronal activity spanned from 50ms before the Go signal up to 50 ms after movement onset) (see Figs. 4-6). The correct-stop trials were grouped based on the length of the SSDs and the neuronal activity was shown from 50 ms before the Go signal. The group number of correct-stop trials varied from 3 to 5 depending on the condition. Three-dimensional trajectories were then obtained from grouped trials by averaging in each group the population activity, eventually smoothing it with a Gaussian kernel (50 ms).

Starting from these grouped spike densities, in the second stage of the analysis we performed the PCA on the concatenated matrix of such average population activities with dimensions N (neurons) x Ctg (conditions x time bins per trial group x number of groups). In each monkey we estimated the Embedding Dimensions from fraction of variance *p*_*k*_ = *λ*_*k*_/ ∑_*j*_ *λ*_*j*_ explained by each principal component (*λ*_*k*_ are the eigenvalues of the covariance matrix) of the neuronal activity. The Effective Dimension *D* = *e*^(*H* − 1) where *H* = − ∑_*k*_ *p*_*k*_ log log *p*_*k*_ is the Shannon entropy. Under the hypothesis of an exponential distribution of the variance, *p_k* ∝ *e*^(−*k*/*D*) provided that *D* ≪ *N*, where *N* is the number single units we considered (51). In all cases the Embedding Dimensions were below 3.

As illustrated in the main text (Fig. 4-6), we further singled out some relevant subspaces. Firstly, we looked for a plane where correct-stop trajectories for a given condition resided. This was defined as the holding plane (Fig. 3 B) best fitting (Singular Value Decomposition; SVD) the cloud of points composing all the three-dimensional trajectories associated to the average population activity for each group of correct-stop trials. We then looked for the axis which best fitted the trajectories of all group of trials projected on the holding plane in the time window between 0 and 300 ms from the Go signal. This holding-and-planning axis (HPA) is represented by a dashed plum dotted line in Fig. 3B, and we found it to be the ideal subspace to represent the first phase of motor plan maturation (always occurring after 200 ms from the Go signal for all conditions and animals), as shown Fig. 4 A. Finally, we defined the planning-and-execution axis (PEA; brown dotted line in Fig 3B, bottom) as the one orthogonal to the holding plane. The trajectory projections onto this direction optimally highlighted the maturation process of the motor plan and the neuronal correlate of the movement execution. The time course of such projections for each trial group is shown in Fig. 4-bottom. The three-dimensional average trajectories for the no-stop, correct-stop and wrong-stop trials in Fig. 3B-botton are shown on a plane after a suited rotation around the HPA.

We also performed another analysis to establish in which of the two subspaces the relationship between neuronal projections and RTs started first. To this aim we considered two samples of data for each subspace, movement direction and monkey: one composed by the times of all the points crossing the thresholds (exclusively for significant regressions), the other composed by the times of the deciles from 4th to 6th. We then compared, for each direction and monkey, the corresponding distributions of time points corresponding to the crossing of the threshold relative to the PEA and to the HPA (Wilcoxon rank-sum test, *P* < 0.01). In both cases the significant relationship started first in the HPA in all conditions considered. In the main text we report the results from the analysis performed on the deciles from the 4th to the 6th, to have an estimate of latency corresponding to the central part of the distributions.

Furthermore, we evaluated the contribution of each unit to the activity in the HPA and in the PEA by calculating a Contrast Index (CI) = (HPA_weight_ − PEAweight) / (HPA_weight_ + PEA_weight_) based on the “weight” of the contribution of each unit to the corresponding axis. We performed this calculation separately for each direction. We then averaged across directions the CIs and ordered the units based on this value.

### Electromyographic recordings and analysis

Electromyographic (EMG) signal was obtained by needle electrodes (monopolar derivation) inserted into the target muscles with the monkey calm. Muscles recorded were: pectoralis, deltoid (anterior, middle, posterior), infraspinatus, biceps, triceps (Fig. S5 A).

Signal from each trial was sampled at 3052 Hz, then band-pass filtered (60-1000 Hz), rectified and down sampled to 1000 Hz. Finally, it was smoothed by using a moving window of 30 ms (in 1 ms steps) and averaged across trials. Data reported are from 6 recording sessions. For further details, see Supplemental Material and Methods.

## Author contributions

P.P. and S.F. designed research; P.P, E.B., V.M., M.G, F.G. performed research; P.P., M.M. analyzed data; P.P., S.F., M.M., wrote the paper. All authors revised the work critically.

## Acknowledgments

In part funded by EU H2020 Research and Innovation Programme, Grant 945539 (HBP SGA3) to SF and to MM, and by Sapienza H2020 2017 grant to SF.

## Supplementary Results

### Time of significant contribution to movement inhibition at the single neuron level in PMd

We focused on conditions with negative time lags between neuronal modulations reflecting movement inhibition (differences in activity between correct-stop trials and latency matched trials) and the corresponding SSRT (see Supplementary Material and Methods; 148/278 [2 movement directions × 139 units]), i.e., the conditions for which the neuronal modulation was possibly causally related to inhibitory behavior. After excluding statistical differences between monkeys [F(1, 146) = 0.24; p = 0.63; monkey P: -60.4 ± 4.7 ms, 95% CI = {-69.7, -51.4}; monkey C: -64.3 ± 6.5 ms, 95% CI= {-77.7, -52}], the population average of the time lags across all monkeys resulted -61.0 ± 3.8 ms, 95% CI={-69.5, -54.1} ms (Fig. S4).

We also found that within the state space approach the neuronal modulation signaling movement suppression at the population level could be detected slightly before the one estimated from the analysis of the single unit activities: Monkey P = -103 ms (CI: -126, -80) vs -60.4 ms (CI: -69.7, -51.4); Monkey C = -76 ms (CI: -84, -67) vs -64.3 ms (CI= -77.7, -52). Overall, these results show that a subpopulation of task modulated units in PMd is capable of signaling the inhibitory reaction needed to stop a movement starting already around -80/100 ms before the end of SSRT.

Importantly, when the population starts to encode the movement suppression, muscles are not activated yet (or, occasionally, just slightly activated, and then rapidly suppressed; see Supplementary Results - below - and Fig. S5).

### Weak neuronal modulations are observed after Stop signal in wrong-stop trials

Once we found that PMd units selectively change their activity in the proper time (i.e., before the end of the SSRT) to inhibit a movement, we also asked whether the Stop signal was capable to affect the neuronal activity in wrong-stop trials, i.e., when the movement was generated irrespective of the command to halt. Observation of neuronal activity in wrong-stop and latency-matched no-stop trials shows that no difference is detected at the moment of movement generation (see unit examples in Fig. 1C, bottom row).

To this aim we considered all the 148 conditions indicated above, and we compared Contrast Indexes obtained from correct-stop vs latency matched no-stop trials to the corresponding Contrast Indexes obtained from wrong-stop vs latency matched no-stop trials. (Fig. S2, Correct vs Wrong, see Supplementary Materials and Methods for details). Positive Contrast Indexes indicate a higher activity in no-stop compared to stop-trials, and vice versa.

If in wrong-stop trials the Stop signal can affect neuronal activity, we should observe non-null Contrast Indexes and their values should be close to those observed in comparisons regarding correct-stop trials. Values either close to zero or different from the Contrast indexes in correct-stop trials represent low or null effects of the Stop signal in wrong-stop trials.

We ran a mixed ANOVA with factors stop trial (correct- or wrong-) and sign of the Contrast Index (either Positive, i.e., higher activity in no-stop trials, or Negative, corresponding to lower activity in no-stop trials). We found that both Positive and Negative indexes in correct-stop trials were significantly different from the ones in wrong-stop trials (Negative Correct: -0.50 ± 0.03, 95% CI = {-0.57, -0.43}; Negative Wrong: -0.11± 0.05, 95% CI= {-0.19, -0.02}, p < 10^-5^; Positive Correct: 0.48 ± 0.01, 95% CI= {0.45, 0.51}; Positive Wrong: -0.050 ± 0.019, 95% CI= {-0.09, -0.012}, p < 10^-5^; F(1, 236)=239, p < 10^-4^. Newman-Keuls post-hoc test, MSE = 0.061, df = 470).

These data show that even if there is some modulation during wrong-stop trials, overall it is weaker compared to correct-stop trials. Indeed, despite an attempt to inhibit the movement, the neuronal activity in wrong-stop trials is not different from the neuronal activity in no-stop trials. This can be also observed when looking at the neuronal activity preceding the movement onset (see Fig. 1C, in the main text, for single unit examples).

### In the specific task, muscles are weakly activated in stop trials and premotor units signal movement inhibition before muscles

Which is the role of the stop-related activity? Is it associated with muscle activations accompanying movement inhibition? Or alternatively, does it aim at interrupting the transformation of the motor plan into an overt action?

To address these questions, we recorded the electromyographic (EMG) activity of the arm muscles participating to the reaching movements (see Fig. S5A), as done in previous studies (27, 52, 53). These sessions were different from those of neuronal recordings. However, the behavior of the EMG sessions conformed to the race model: the wrong-stop trials were faster than in no-stop trials (average difference across sessions = 38 ± 12 ms, t-test (5) = -6.6, p = 0.0012); most importantly the SSRT estimates (mean ± SD = 192 ± 25 ms) were very close to the SSRT estimates obtained in the neuronal recording sessions from the same monkeys (see Table S1).

Most of the time, when movements were successfully inhibited in both directions, the EMG activity did not show any significant modulation (for an example muscle see Fig. S5B). This shows that, at least in the context of this task and setup, stopping an arm movement may not require any muscle activation. More specifically, we found that in the large majority (82; 66%) of all conditions (124: direction × muscle × session, criterion of at least 5 trials per condition) there was not any significant muscle modulation in correct-stop trials in the 300 ms-window following the Stop signal. For the other 42 conditions, we compared the EMG activity of no-stop trials and correct-stop trials to extract the latency of the first significant difference. In the conditions for which this latency could be estimated [38 out of 42, 90%; in 4 conditions the difference did not reach the criteria (see Supplementary Material and Methods)], it always preceded the behavioral SSRT. In this subset, we obtained the estimate of the electromyographic time lags by subtracting the latency of the first significant EMG difference from the behavioral SSRT. This is the electromyographic counterpart of the neuronal time lag. Muscle modulation related to inhibition followed the neuronal modulation: the muscular time lag indeed was, in absolute value, smaller than the neuronal time lag: -38.2 ± 7.1 ms, 95% CI = {-52.3, -24.1}; (F(1, 184)=8.6, *p* = 0.004), see Fig. S5C. Thus, the EMG signature of the stop process was either an unmodified level of activity, or when a small increase of activity was observed this was suppressed always after inhibition related neuronal modulation, and before behavioral SSRT suggesting that a proper action cancellation requires a control before muscle activation, at least in the version of the task we used.

We confirmed that the muscles we recorded from, were involved in movement generation by comparing their patterns of activations between opposite directions: all the muscles were active before movement generation and showed directional dependence (Fig. S5 D-G). However, although the muscles showed directionality, they did not show clear agonistic or antagonistic activations when compared between opposite movement directions. A possible reason for this is that the arms of the monkeys - having to act on stimuli presented on the touch screen - had to contrast the force of gravity throughout the phases of the trial, so the muscles had to be always active at some level. This experimental context is different from the ones of other studies where the use of a cursor or a joystick in a 2-D plane probably helped in generating agonistic- antagonistic patterns (for example, see 49).

We also wanted to investigate the differences in muscle activation between wrong-stop trials, and correct-stop trials, aligned to the Stop signal. We expected that in wrong-stop trials the muscle activation must occur before the end of the hypothetical SSRT to start the movement. We tested this hypothesis by calculating the latency of the difference between wrong-stop and correct-stop trials (Fig. S5F and G). From a sample of 36 conditions (at least 7 trials for correct- and wrong-stop trials, with the same SSD) we found that the latency of this difference was 74 ± 40 ms (mean ± SD) from the Stop signal (maximum detected latency was of 134 ms). Thus, well before the end of the behavioral SSRT (more than 100 ms) muscle activity in wrong-stop trials was already too high to be suppressed.

In wrong-stop trials the monkeys typically started the reaching movement and then halted it before they could reach the peripheral target. Based on this observation we asked whether and how the presentation of the Stop signal might affect muscle activity during the movement phase of the wrong-stop trials. We then compared the muscle activity between the wrong-stop trials and latency matched no-stop trials. Indeed, the first phase of the movement is the same between the two trials (see Fig. S5H for an example muscle), while the differences emerge only once the Stop signal is encoded in wrong-stop trials and the reaching movement is then halted. To increase the power of our analysis, we selected no-stop trials in the range of the RTs ± interquartile range (IQRT) observed for wrong-stop trials (for wrong-stop trials we considered conditions with at least 6 wrong-stop trials per SSD). Considering all 88 conditions (SSD × direction × muscles) for which the comparison was possible, we found differences between the two types of trials in the 150 ms preceding RT (comparisons between average activities in windows of 40 ms and 1 ms step, of wrong-stop and no-stop latency matched trials) only in 2 conditions. We typically observed that a clear difference between wrong-stop and no- stop trials occurred after the movement started, purportedly related to the difference in movement between the two trial types. We decided to calculate the latency of this difference relative to the movement onset, and we found a significant difference for 69/88 conditions; on average this value was 71 ± 43 ms (Fig. S5I). In summary, about 70 ms after the movement onset a clear change in EMG activity probably reflected the action of the online movement stopping process in wrong-stop trials.

On average we estimated that EMG activity started to increase above baseline about -82.5 (71.9) ms; 95% CI = {-104, -61} relative to movement onset (values obtained across 48 conditions used to estimate this value; threshold was set to average baseline + 3SD).

In agreement with Logan (54) an incoming movement can be stopped only if the stop process acts before the point of no return, i.e., the point when neuronal and muscle activities supporting movement generation are too advanced to be suppressed by it. This last case is observed in wrong-stop trials, in which the Stop signal is presented when the movement planning-execution is too advanced, and thus the movement is generated despite its presentation. By applying the race model, we can consider the end of SSRT as coincident to the movement onset (54). In this view, to be effective the Stop signal must be presented around 190 ms before the movement starts. In this study we found that muscle activity starts to rise about 85ms before movement onset and that the modulation of muscle activity following a Stop-signal can be detected in some cases around 40ms before the end of SSRT. Thus, neuronal activity must either impede the rise of or suppress a low level of muscle activity before and not later than 85-40 ms before movement onset to yield an effective inhibition We found that the latency of neuronal population signal reflecting suppression can start already around 80 ms before the end of SSRT, suggesting that the information to stop arrives on average when the EMG activity is - possibly - only beginning to rise, as confirmed by our observations on EMG activity.

### HPA activity reflects movement planning

To better understand the role of the HPA activity in movement generation we implemented a **Context informed Go-NoGo task**. This task was characterized by the following sequence of stimuli and events: at each trial a visual Cue defining the context (Cue-Context-signal, see below) was presented for 200-300 ms as soon as the monkey touched the central target. Then a Target appeared either to the left or to the right. After a variable delay (800-1200 ms) a Command Signal appeared, instructing the monkey either to reach towards the Target (Go signal), or to keep the hand in position (No-Go signal).

The Cue-Context signal allowed us to modulate the activity of movement planning. Indeed, 3 different Contexts where defined: the **Advanced Go** (see the Fig. S11), informed the monkey that at the end of the trial the Command Signal was a Go signal; the **Advanced NoGo** informed the monkey that at the end of the trial the Command signal was a NoGo signal; and the **Go-NoGo** context that informed the monkey that in 50% of the trials a Go signal would appear as Command signal, while in the others a NoGo signal would.

Considering all the trials, 37% were Advanced Go, 21% Advanced NoGo, 21% Go trials, 21% NoGo trials. Thus, four classes of trials were collected: Advanced Go in which the monkey was sure since the beginning of the trial that a movement was required; Advanced NoGo in which the monkey knew since the beginning of the trial that no movement was required (and, as such, no planning was required); Go, in which the monkey decided to move after the Command signal; NoGo in which the monkey decided not to move after the NoGo signal.

Using this task, we extracted data from 113 units. Trials were ordered by RT and grouped in deciles and tertiles for Advanced Go and Go classes. For Advanced NoGO and NoGo were ordered by delay durations and grouped in tertiles. We performed the same analysis done on the countermanding (CMT) task on this GoNoGo task (see methods in the main text). In this case the holding space was estimated by considering the activity in the NoGo trials, because they required movement preparation but the Command Signal was a NoGo signal. Somehow, this condition can be considered as a Stop condition with SSD=0. The monkey did not make errors in this condition.

The Fig. S11 shows the activity projection on PEA and HPA of the different trial classes. In the PEA (Fig. S11 A), only activities associated with movement generation (Advanced Go and Go; gray and black traces respectively) emerged above the dotted line representing a threshold level. Interestingly this activity starts even before the Command Signal presentation for the Advanced Go trials. In the HPA all the classes but one (the Advanced NoGo; red traces), show an increase in activity that overlaps up to 200ms following the Command Signal. These classes are associated with movement preparation and planning, because a Command signal requiring movement is certain or probable. Their activities are similar to the ones observed in the CMT for all the trial types (no-stop, wrong-stop, and correct-stop). This is because in all these instances the generation of a movement was possible. In this new task only the Advanced NoGo shows a constant low level of activity in the HPA. Indeed, the cue presentation makes clear to the monkey that no movement will be required. As such, we can hypothesize that the HPA activity is associated with the first stage of movement planning. As such it is necessary for movement generation (because it is always present before movement generation or in cases when a movement is possible), but its presence does not define whether there will be a movement. The generation of the movement occurs only when activity increases in the PEA. This is further supported by the linear relationship between RTs and Advanced Go (gray) and Go (black) trials (Fig. S11 B), similar to the one observed for the CMT task (see Fig. 7 in the main text).

## Supplementary Figures and Tables

**Figure S1.**
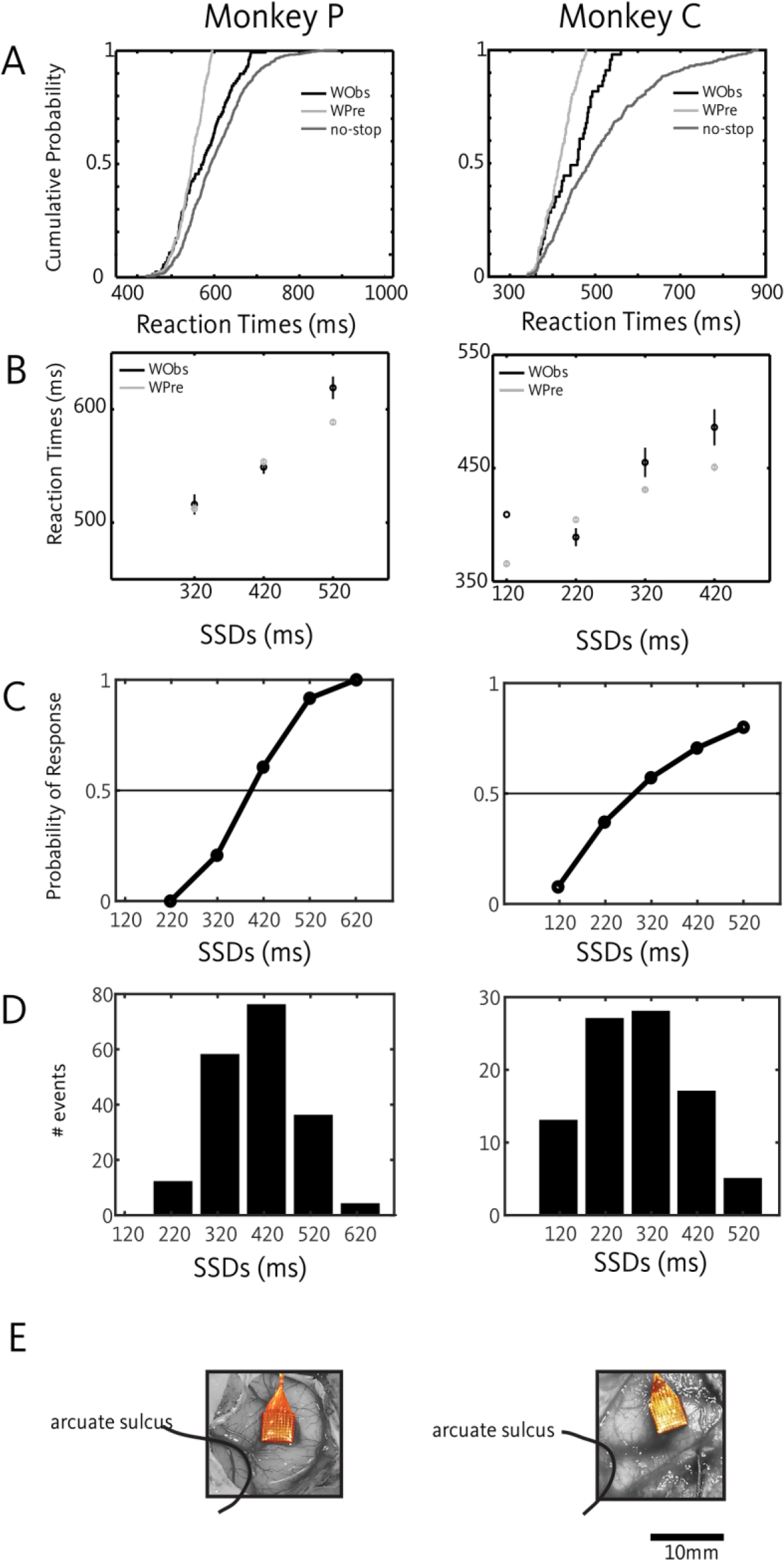
Behavioral performance in the countermanding task and recording sites. **A.** Validation of the race model: for each monkey no-stop, observed wrong-stop (WObs), and predicted wrong-stop (WPre) cumulative RTs distributions are plotted (top row). **B.** The same data are also shown as functions of the SSDs (bottom row). C. Inhibition functions: probability of responding to a Stop-signal as a function of SSDs. D. Distribution of SSDs presented by employing the staircase procedure. **E.** Pictures of the array’s location for monkey P and C.

**Figure S2.**
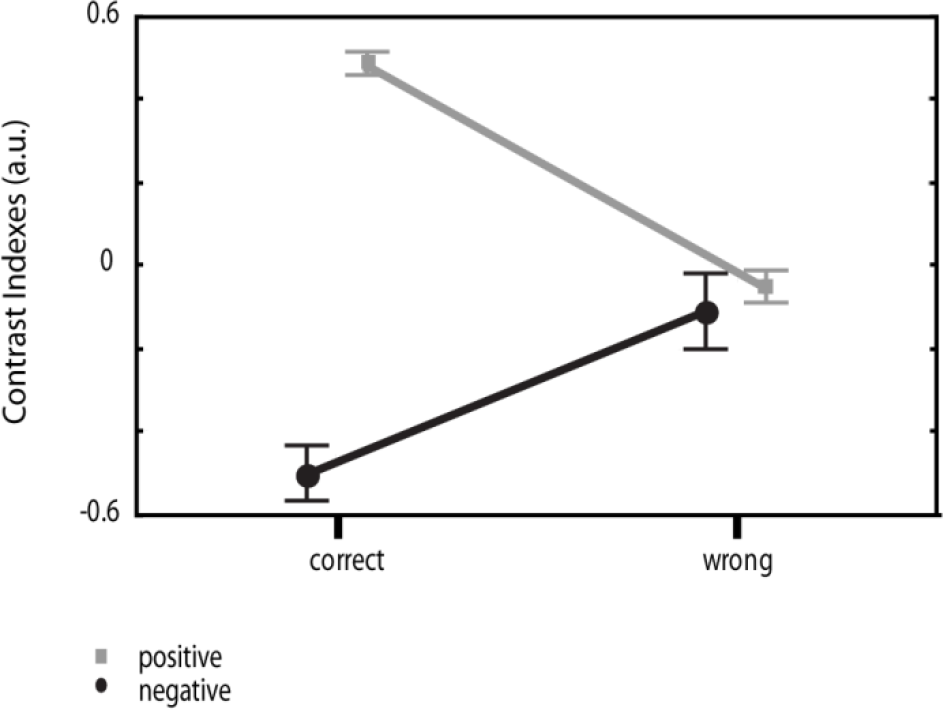
Values of contrast indexes for inhibited (correct-stop trials) vs not-inhibited (wrong-stop trials) movements. Error bars, 95% confidence interval. (see Supplementary Material and Methods for further details).

**Figure S3.**
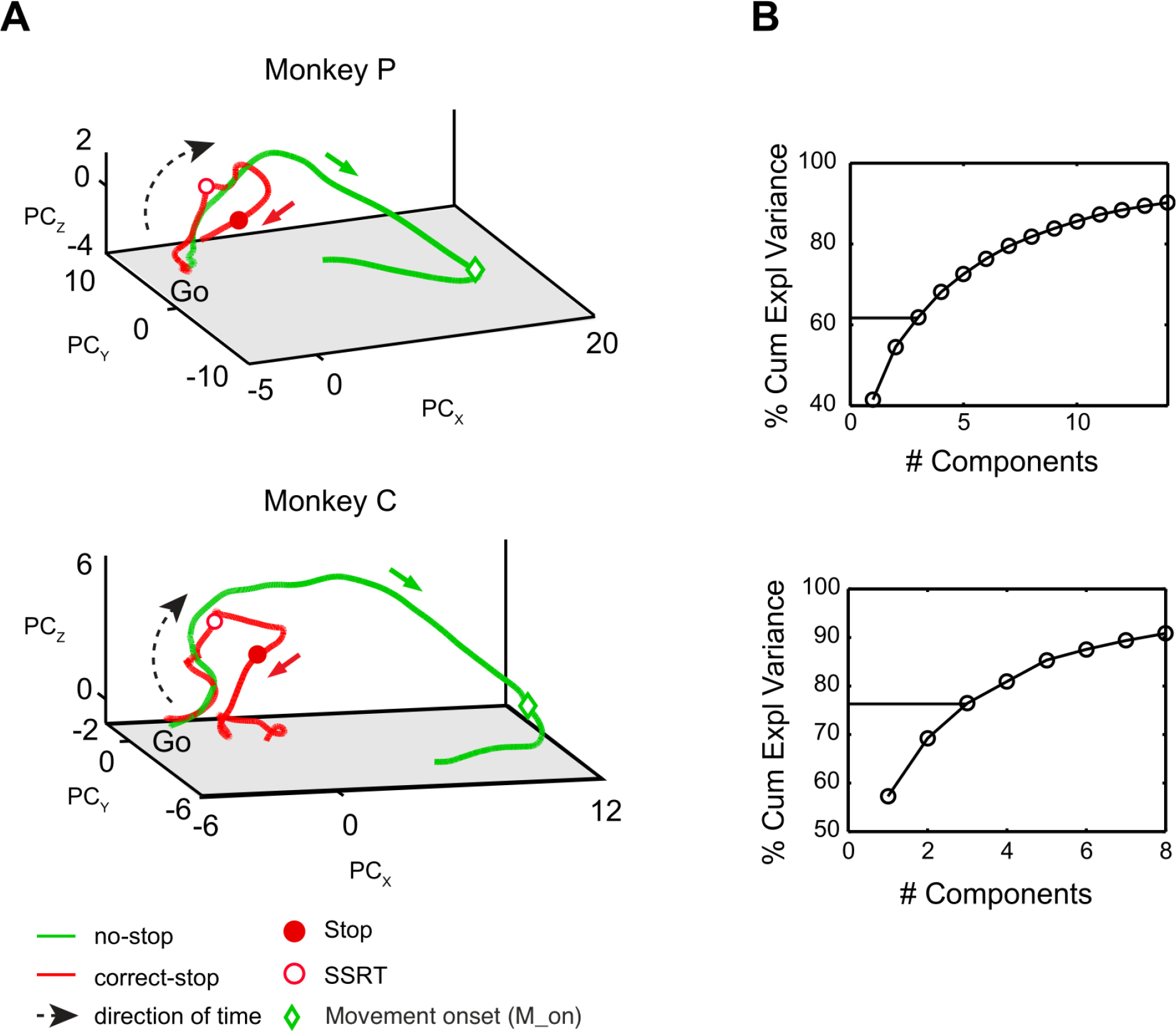
Neuronal dynamics of movement inhibition. Direction 2. **A.** Neuronal trajectories in the state space defined by the first three principal components associated to direction 2 as in Fig. 2A of the main text. **B.** Cumulative explained variance by the principal components capable of explaining up to 90%. The horizontal line delimits the first three components used to plot the neuronal trajectories. Across all monkeys, the first 3 components could explain at least 60% of the variance.

**Figure S4.**
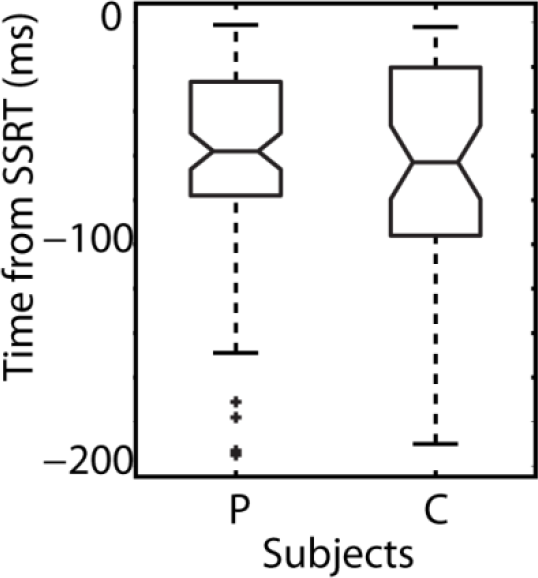
Neuronal modulation occurs before behavioral SSRT. Boxplots illustrate the timing of neuronal modulations (obtained from single unit activities) with respect to the behavioral SSRT for both monkeys.

**Figure S5.**
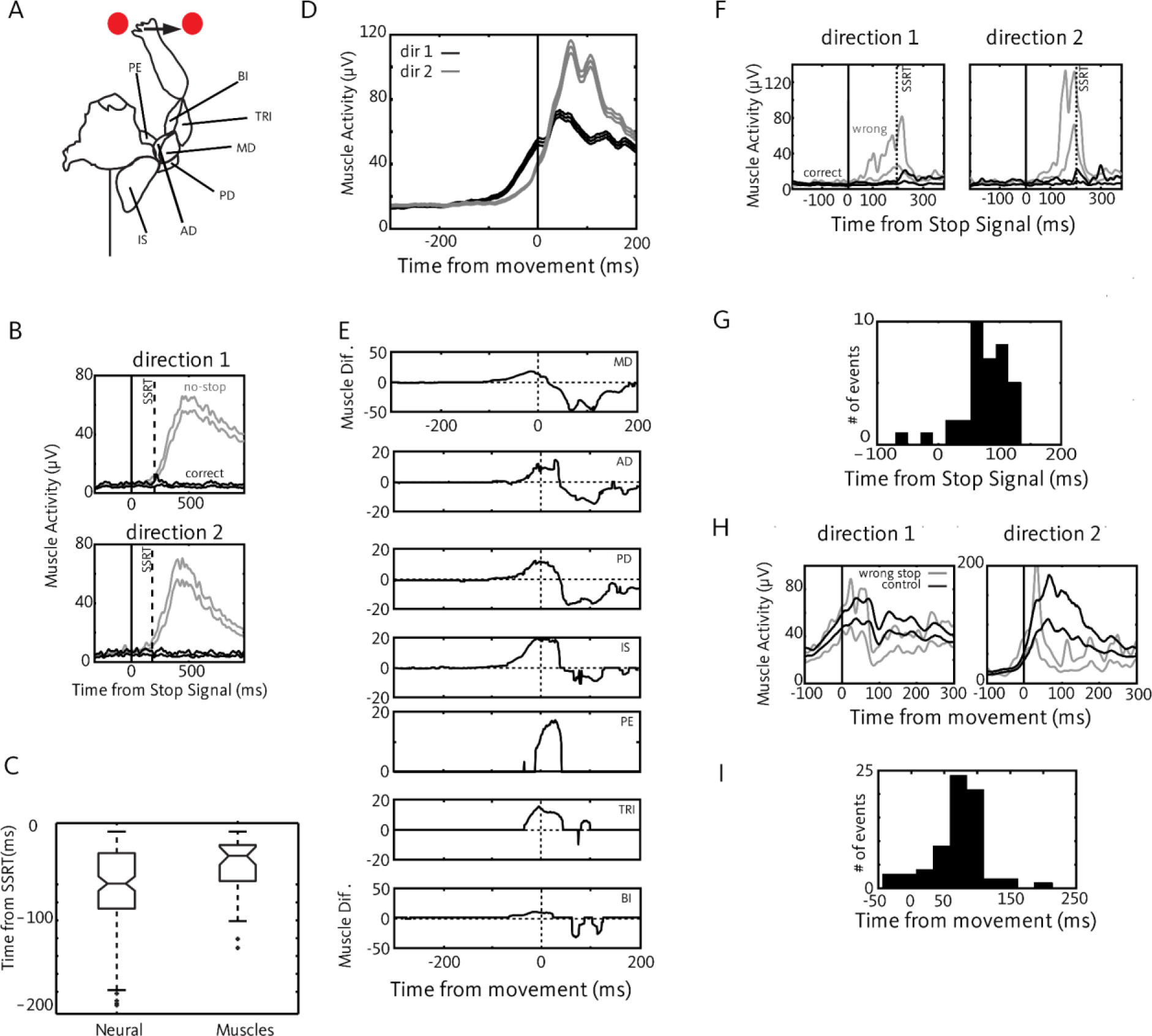
Muscle modulation in the task. **A**. Recorded muscles and schematic of the limb postures during the task. Six muscles were recorded: infraspinatus (IS), anterior deltoid (AD), middle deltoid (MD), posterior deltoid (PD), biceps (BI), triceps (TRI) and pectoralis (PE). **B.** Comparison between muscle activities in correct-stop and corresponding latency matched no-stop trials for the infraspinatus in one experimental session (Monkey C). Each panel represents muscle activity for a single movement direction (mean ± SE). **C.** Boxplots of time lags between stop-related neuronal modulations and SSRT (Neural), and time lags between muscle stop-related neuronal modulations and SSRT (Muscles) for both monkeys. **D**. Average activity for the MD across different sessions (n=4; mean (SE)) and directions. **E**. Activity differences between the two movement directions aligned to movement onset for all the muscles recorded across sessions. Values different from 0 represent significant differences (see Supplementary Materials and Methods). **F**. Comparison between correct-stop and wrong-stop trials for the same SSD in the two movement directions (sess. #3, medium deltoid). **G**. Distribution of the latencies of EMG divergences between correct-stop and wrong-stop trials relative to stop signal presentation. **H**. Comparison of an example muscle activity between wrong-stop and control no-stop trials (posterior deltoid, one session), aligned to movement onset. **I**. Distribution of the latencies of the onset of the EMG differences between wrong-stop and no-stop trials after movement onset.

**Figure S6.**
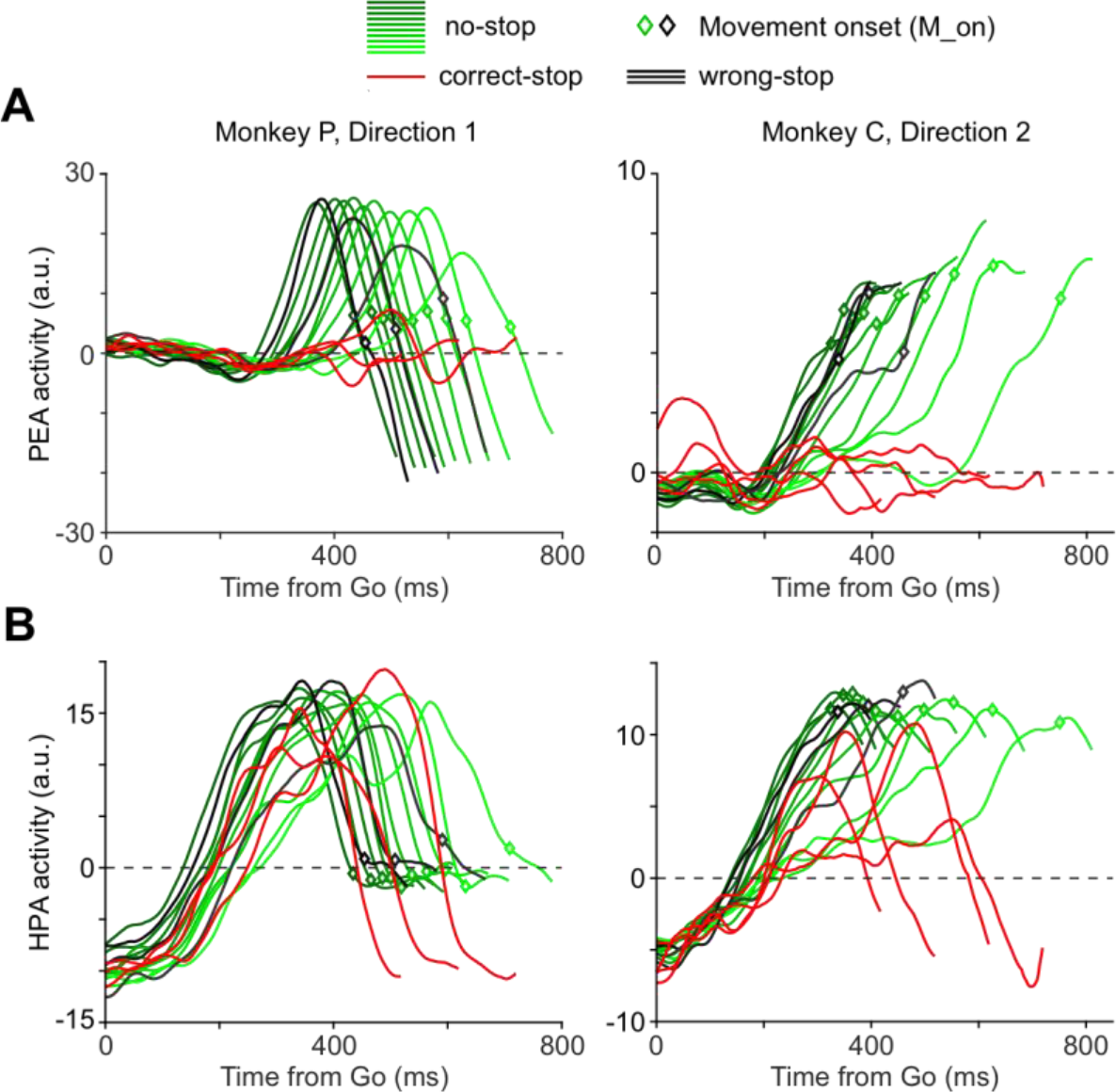
Projections of neuronal activities on the HPA and PEA, Direction 2. **A.** Projections on the planning-and-execution axis (PEA) of the neuronal trajectories (for direction 2) grouped by RTs (no-stop in deciles and wrong-stop trials in third intervals) or SSDs (correct-stop trials), for each monkey. **B.** Projections on the holding-and-planning axis (HPA) of the same neuronal trajectories as above. This figure refers to Fig. 4 in the main text.

**Figure S7.**
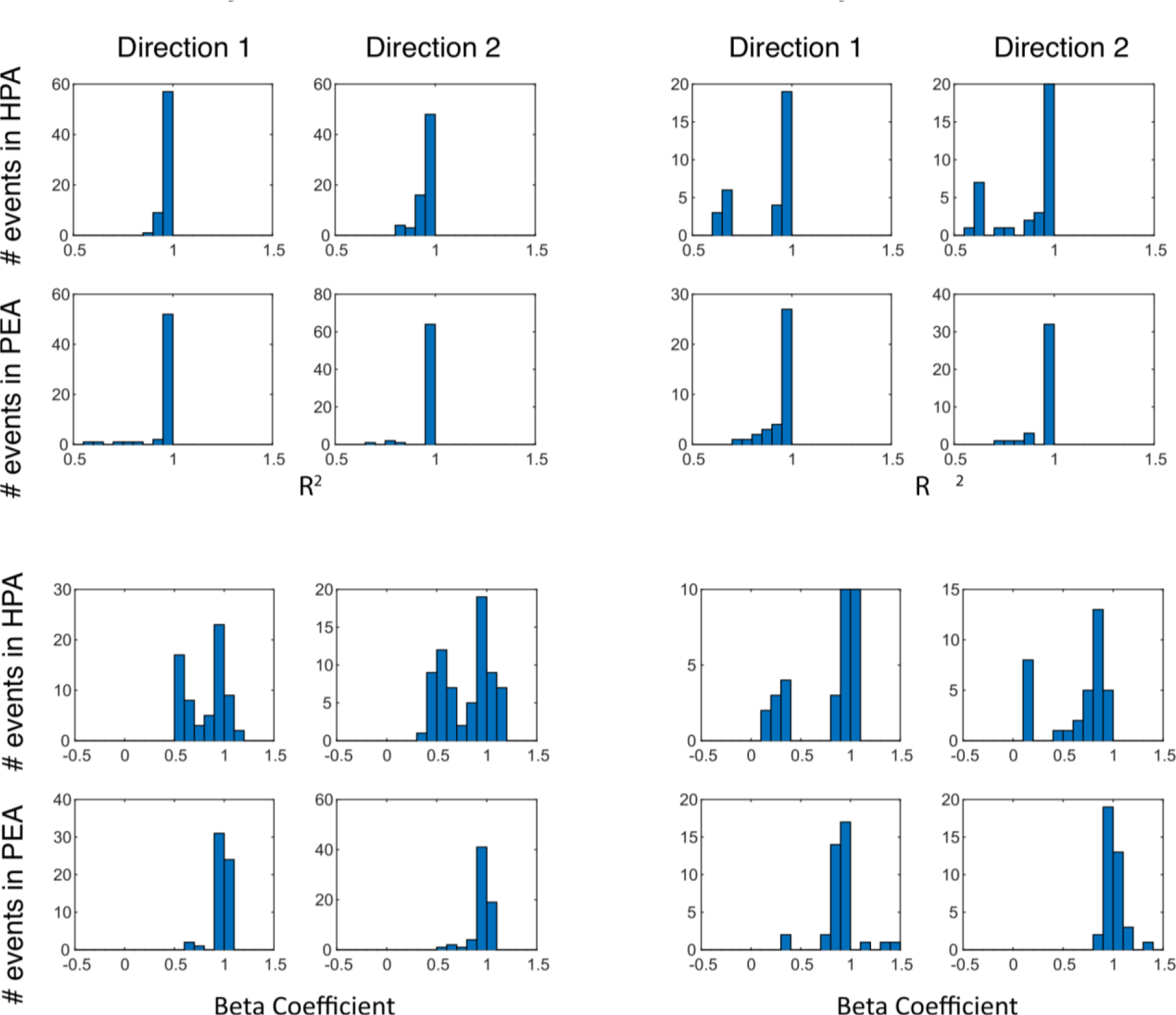
Relationship between PEA and HPA levels of activity, and RTs. For each monkey and direction of movement values of R^2^ and Beta coefficient of the regression analysis between the values extracted by using the thresholding method (see Supplementary Materials and Methods) and RTs are reported.

**Figure S8.**
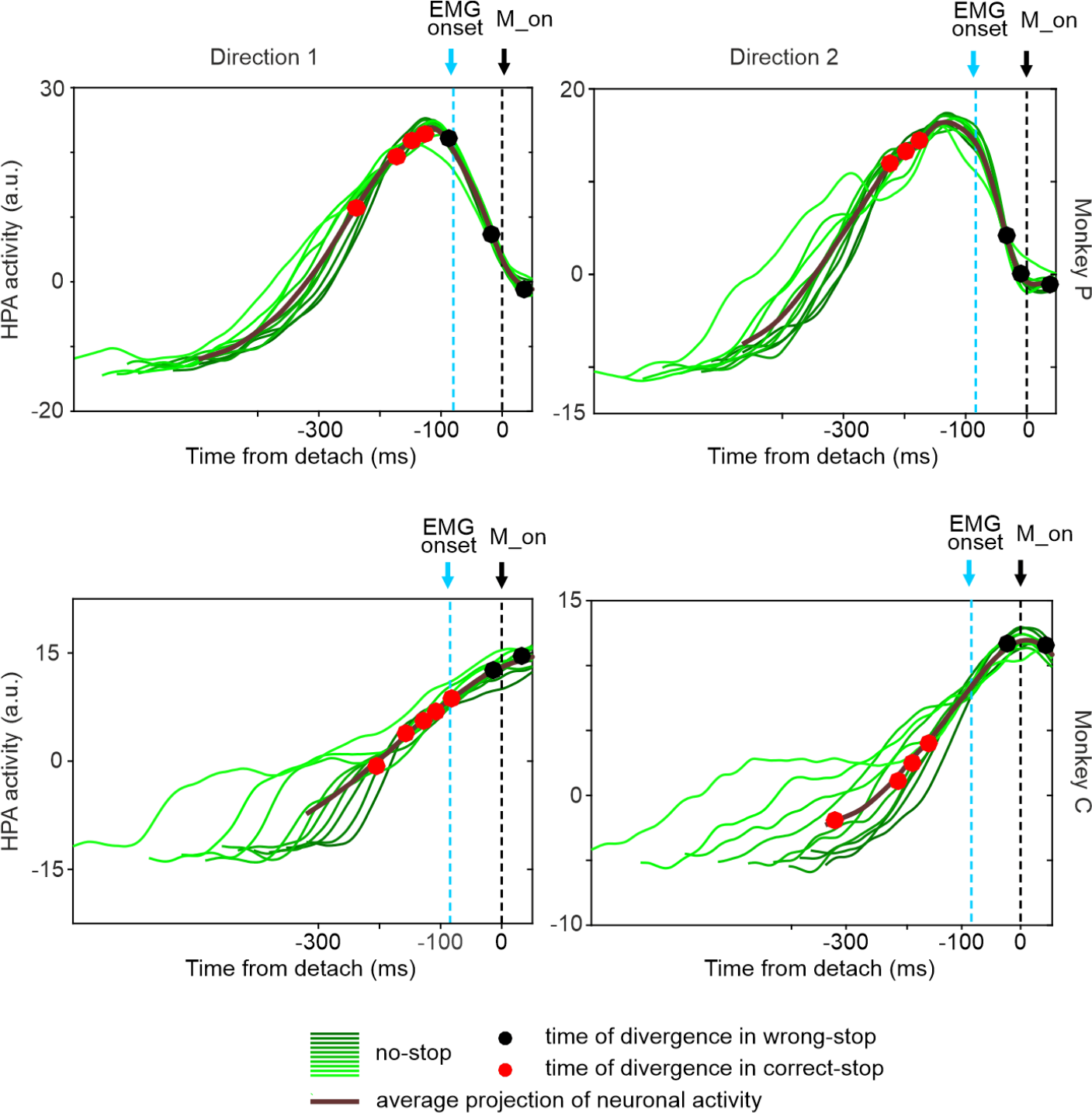
HPA no-stop activities grouped by RTs and aligned to movement onset. Data from both animals. The solid line represents the average of the activities (also sorted by deciles and in grey levels) projected in the HPA. Red (correct-stop) and black (wrong-stop) dots highlight the level of activity observed at the time of divergence. For correct-stop trials this activity corresponds to the level of activity observed at the time of divergence between correct-stop and no-stop trials. For wrong-stop trials we employed the level of activities identified in Fig. S6 and corresponding to the average time of divergence. In correct-stop trials the modulation occurs before muscle activation; in wrong-stop trials the modulation occurs too late to halt the movement.

**Figure S9.**
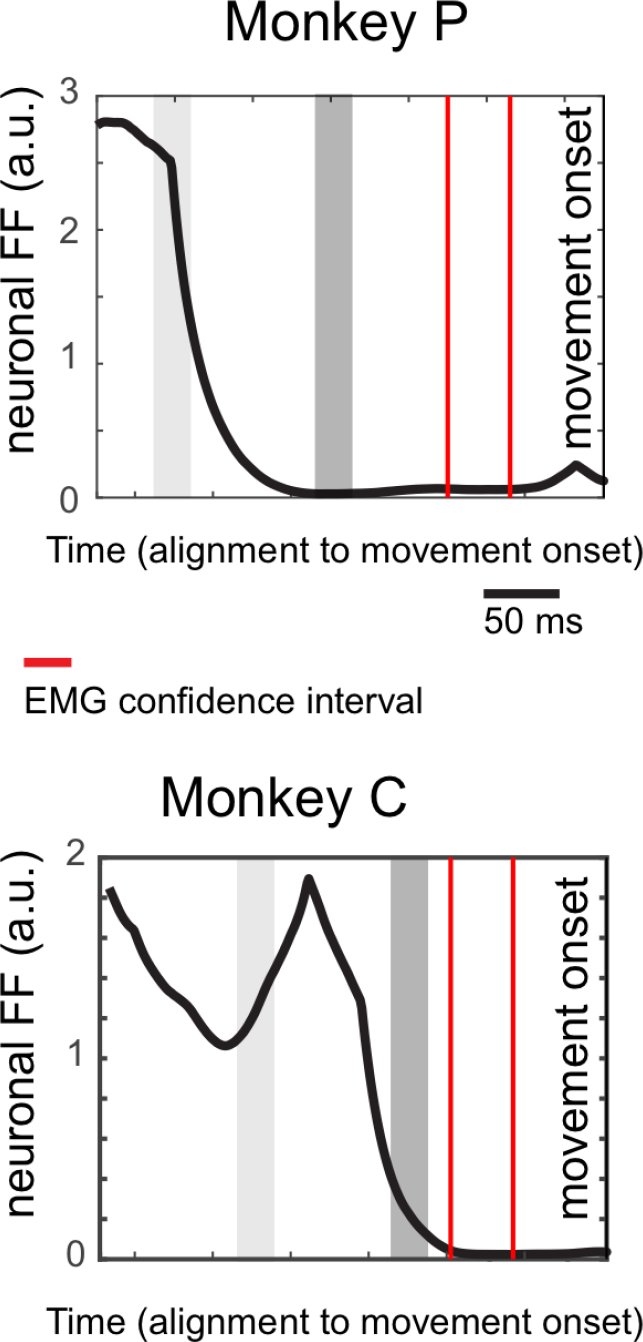
Reduction of variance of the HPA no-stop activity before movement onset. The neuronal variability (Fano Factor; FF) is computed every each millisecond, as the variance of the HPA activity across trials grouped in deciles (for both movement directions) and divided by the mean value. Gray regions represent epochs (40 ms interval) where values of high (light gray) and low neuronal variability (dark gray) have been measured and used for data in Fig. 5. The temporal window indicated by the two vertical redlines is the EMG onset confidence interval.

**Figure S10.**
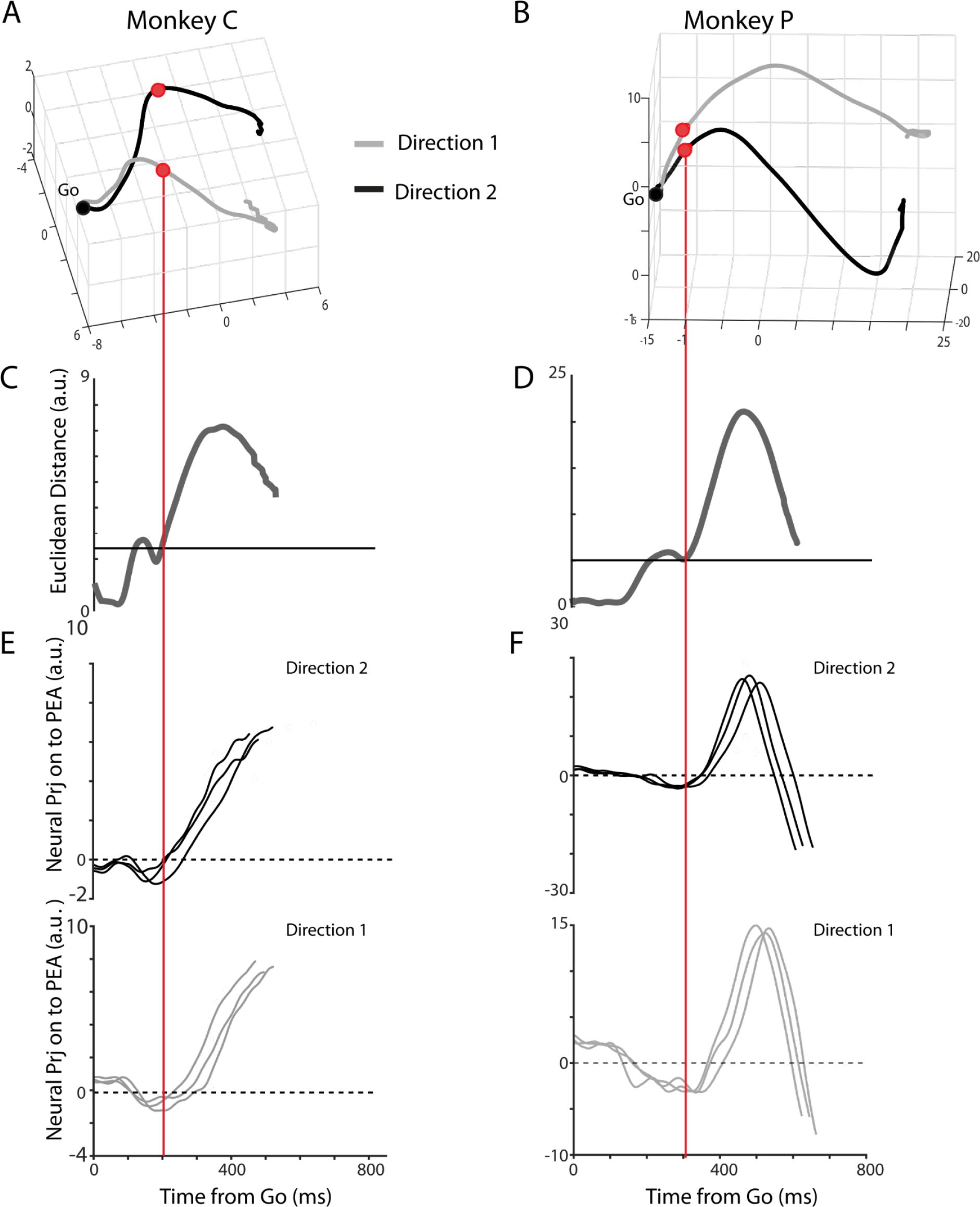
Directional information is reflected in neuronal trajectories. **A** and **B**: average neuronal trajectories of no-stop trials for the two movement directions in the 3D subspace of the first three PCs. **C** and **D**: Euclidean distances between trajectories for different movement directions: at latency of about 200 ms (300 ms) from the Go signal for monkey C (P) the two trajectories strongly diverged (the horizontal line represents a threshold computed as the mean ± 3SD of the Euclidean distance in the epoch from -50 to +50 ms around the Go signal). **E** and **F**: neuronal activities in the PEA corresponding to the 4^th-,^ 5^th^ and 6^th^ decile of RT, separately for each direction.

**Figure S11:**
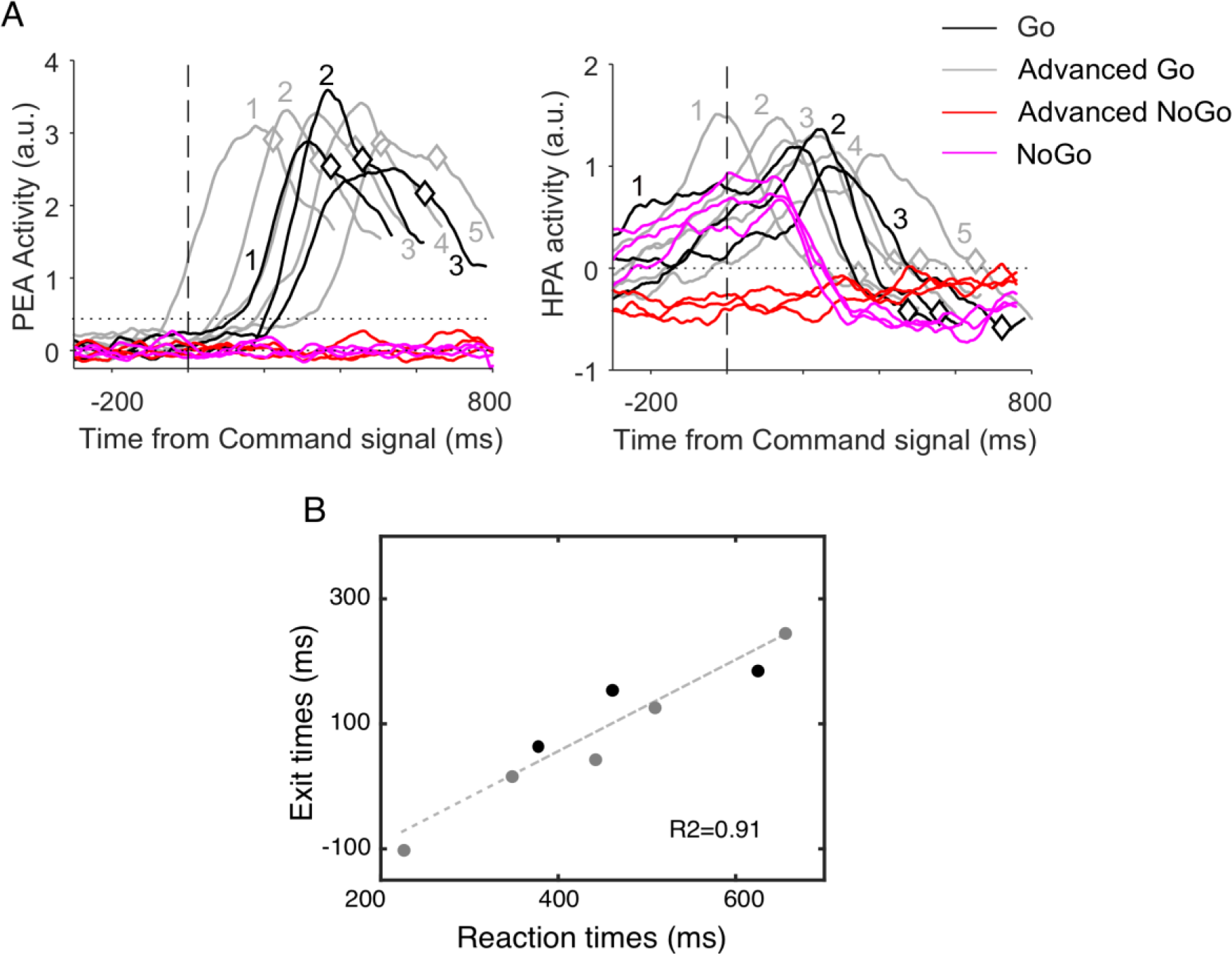
**A**, PEA and HPA dynamics in the Context informed Go-NoGo task, respectively. **B**, relationship between exit times (crossing between the dotted line in PEA 0.5 (a.u.) and activities in Go and Advanced Go trials and corresponding RTs (dark dots: Go trials; gray dots: Advanced Go trials).

**Table S1.**
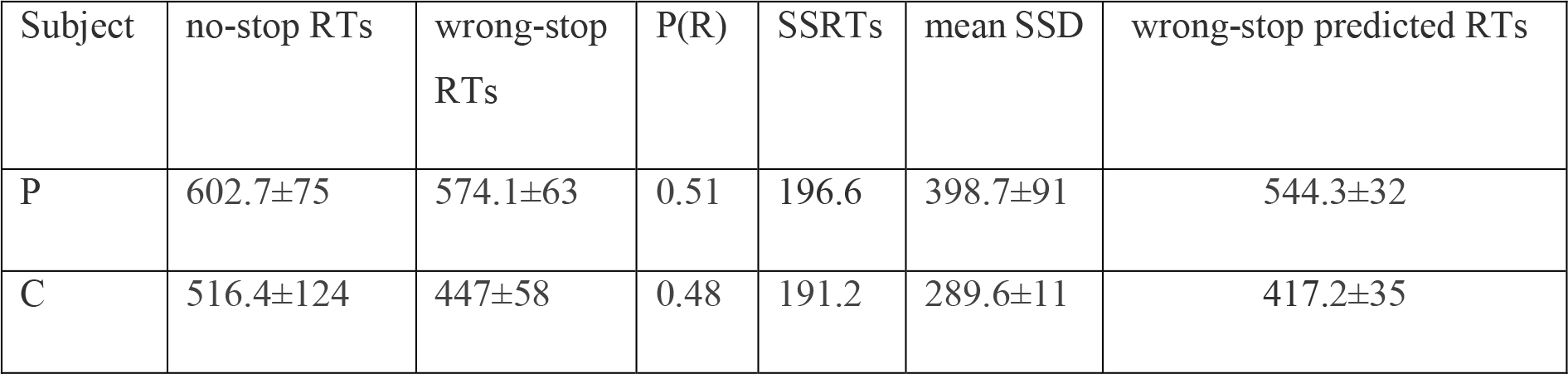
Behavioral results (related to Figure 1). For each monkey the mean ± SD of the behavioral indexes is reported. All values, except the probability of response P(R)) are in milliseconds (ms). Analysis of behavioral data for all the monkeys have shown that the wrong-stop RTs were shorter than RTs in no-stop trials (P < 0.05, Wilcoxon rank-sum test). This is a prerequisite to estimate the SSRT (see also 43).

## Supplementary Material and Methods

### Subjects

Monkeys were pair-housed with cage enrichment (toys, foraging devices). They were daily fed with standard primate chow supplemented with nuts, raisins, and fresh fruits if necessary. The monkeys received their daily water supply during the experiments.

All the surgeries were performed under sterile conditions and veterinary supervision. Antibiotics and analgesics were administered postoperatively. Anesthesia was induced with ketamine (Imalgene, 10 mg kg^− 1^i.m.) and medetomidine hydrochloride (Domitor, 0.04 mg kg^− 1^ i.m. or s.c.) and maintained by inhalation isoflurane (0.5–4%) in oxygen (5 l min−1). Antibiotics were administered prophylactically during surgery and postoperatively for at least 1 week. Postoperative analgesics were given at least twice per day. Recordings started after full recovery from surgery (3/4 weeks).

### Behavioral analysis

To validate the race model’s independence assumption and compute the SSRT, the mean RT of wrong-stop trials must be shorter than the one of no-stop trials (see for example 21, 43). Other tests typically employed to validate such assumption, require that the mean RT in wrong-stop trials should increase with increasing duration of the SSD and that the same mean RT in wrong-stop trials should be equal to those predicted from the race model for each SSD. This second test requires calculating the expected wrong-stop RT from the no- stop trials distribution of RT. For each SSD, a threshold RT given by the sum SSD + SSRT is considered. The distribution of expected RT for wrong-stop trials is then assumed to be the same as the distribution of no-stop trials RT shorter than the above threshold RT. These predicted RTs are typically tested when the SSD does not dynamically change. Indeed, a reliable estimate of expected RTs requires many trials for each SSD (40, 55). Nevertheless, we ran these tests of the independent race model also on our data, although SSD distribution was adjusted depending on the animal performance. We obtained that considering all the values, wrong-stop predicted RTs were shorter than wrong-stop observed RTs for both monkeys (rank-sum test: p<.0001 and p<.002, monkey P and monkey C respectively). No differences were found comparing the mean values of RT for each SSD in Monkey P (p=0.1) while a difference was found for Monkey C (p=0.03). Furthermore, in both cases mean RT in wrong-stop increased with increasing SSDs (Spearman corr =1 for monkey P and Spearman corr =0.8 for monkey C). Overall, the results support the independence assumption (see Figure S1 and Table S1).

### Neuronal analysis

One purpose of the study was to confirm, as previously observed by our group (8), that single neurons in PMd were related to movement inhibition, i.e., if they were differently modulated when the movement was inhibited or when the movement was executed. To this aim we compared the neuronal activity (namely, the spike density function, SDF) in stop trials to the neuronal activity in the latency-matched no-stop trials (also defined as control trials). To perform such comparison, the neuronal activity in the latency-matched no-stop trials for each SSD were aligned to the onset time of the hypothetical Stop signal (i.e., the SSD) working out the SDF. The same procedure to calculate the average activity was used for correct-stop and wrong-stop trials.

We calculated a differential SDF by subtracting the SDF of correct-stop trials from that of latency-matched no-stop trials to derive a stop-related neuronal latency modulation (see Materials and Methods in the main text, for details).

To establish when neuronal modulation occurs relative to the estimated SSRT we subtracted the stop-related neuronal latency from the SSRT estimated in the same session. This allowed us to evaluate the time lag between the behavioral inhibition and its neuronal correlate at the single neuron level.

We also calculated a Contrast Index, by contrasting the average firing rate in the 50 ms following the onset of the stop-related neuronal latency in correct-stop trials to the corresponding 50 ms of average firing rate in the control (latency matched; see main text) no-stop trials, as follows:

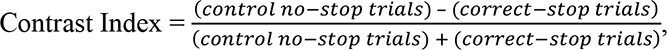

### Using the same approach, a Contrast index was calculated for wrong-stop trials

These Contrast Indexes were used to compare the effect of the Stop signal in inhibited (correct -stop) *vs* not inhibited (wrong-stop) trials. Contrast Indexes were positive (negative) when neuronal activity was higher (lower) in control no-stop trials compared to the corresponding correct-stop and wrong-stop trials. In all neurons with no significant stop-related neuronal latency, the Contrast Index was set equal to 0.

### Electromyographic recordings and analysis

To establish how the muscles recorded participated in arm movements generation and inhibition, we compared their patterns of activation for movements toward the two possible opposite directions (left and right). For each muscle we compared all trials with planned movements towards one direction vs trials towards the other direction aligned to movement onset in 10 ms, not overlapping steps (Wilcoxon rank-sum test, p was set to 0.0001 because of the number of comparisons). From this analysis we obtained a vector of zeros (no difference) and ones (significant difference) to be multiplied by the differential obtained by contrasting the EMG activity of the two directions. Thus, only significant differences had values higher than 0.

To study the muscular modulations that characterize movement inhibition, we selected those conditions with at least 5 correct-stop trials per single SSD. We first established whether in correct-stop trials EMG activity increased respect to the 300 ms preceding the Stop signal (at least 50ms after stop signal presentation above 2.5 SD of the EMG activity in the 300 ms preceding the stop signal). Then we estimated the electromyographic equivalent of the average time lag between the neuronal modulation after the Stop signal presentation and the SSRT. To this end, we first calculated the differential EMG activity between correct-stop and latency matched no-stop trials and we searched for values above a level for at least 50 ms. The baseline level corresponded to the mean + 3 SD of the differential EMG activity in the 400 ms preceding Stop signal presentation). Once the criteria were respected, for each muscle we considered as electromyographic latency the difference between the SSRT and the first time point above the baseline level.

We also calculated the latency of the divergence in EMG activities between wrong-stop and correct-stop trials. To perform this analysis, we considered all SSDs with at least 6 trials of each type. We calculated as baseline the mean difference between wrong-stop and correct-stop trials in the 400 ms around the time of the Go signal presentation. We then considered as latency of the divergence the first time point after the Stop signal presentation in which the difference between the wrong-stop and the correct-stop trials was above the mean ± 2 SD and continued to stay above this level in the following 75 ms.

To establish the latency of the difference in EMG activation between wrong-stop trials and no-stop trials with similar RT, we subtracted EMG activity of no-stop from the one of wrong-stop trials aligned to the Movement onset. The latency was the first time point when the difference surpassed for at least 150ms the limit defined by the average difference in the -300 to -100 ms before Movement onset of at least 2 SD.

### Moving threshold regression analysis between neuronal projections and RTs

We performed regression analyses to have an account of the temporal evolution of neuronal activity after the Go signal and to establish the relationship between this evolution and the RT.

These analyses were conducted separately on the neuronal projections in the PEA and HPA, and the corresponding RTs. Starting from the minimum neuronal activity we lifted a threshold activity up to the highest level (50 steps) of activity reached. For each step we extracted the times when the activity crossed these thresholds, and we regressed these times against the corresponding RTs. This was done only if we were able to obtain at least 7 values of the crossing times. We then calculated the regression slope (Beta coefficient, see Figure S7), and we kept it if the regression analysis was significant (p < 0.01). When the projections showed an oscillation (see for example PEA activity in monkey P) we performed twice the procedure to account separately for two groups of crossing. For the initial part of the PEA (see Figure 4A and the relative description on the main text) we observed a decay of activity drawing a through that did not show any relationship with RTs.

The Beta coefficients observed were in line with previous works (see 15). We expected to find a broader distribution of slopes for activity showing a ramping like activity: indeed, these profiles usually start with regression slopes well below 1 that increases over time; differently it is possible to have stereotyped responses for which regression slopes are very close to 1 throughout most part of the trial.

